# Signals of consistent genetic diversity decline are not yet measurable in global meta-analysis

**DOI:** 10.1101/2025.08.01.663988

**Authors:** Moises Exposito-Alonso

## Abstract

Genetic diversity within species must be conserved as the basis for evolutionary adaptive capacity. To inform new global conservation policy, Shaw et al.^1^ conducted a meta-analysis of temporal genetic diversity data for over 600 wild species spanning a median of 7 years, reporting significant overall diversity loss of effect size g = −0.1. My re-analysis translating this effect to percentage change finds no robust statistical support for consistent decline, with overall changes near zero—from −1% to +1% diversity change depending on the averaging method. When testing each species individually, roughly half increased and half decreased in diversity. While genetic diversity is expected to decline following population losses, evolutionary theory predicts such declines substantially lag behind population reductions, and thus should not be measurable on average at the timescales in Shaw’s data, except in dramatically collapsing species. However, conservation policy should not wait for consistently measurable genetic diversity loss—to avoid future lagging losses, we must protect populations now.

---

Global biodiversity—from ecosystems, species, and genetic diversity within species—is in decline^2^. Genetic diversity enables species’ long-term evolutionary adaptation and survival ^3^. To inform conservation priorities ^4^, we need to quantify diversity declines, but measuring genetic diversity loss has been challenging ^5–9^. Shaw et al. (ref. ^1^, hereafter Shaw) compiled a database of 628 species with temporal samples and conducted an effect size meta-analysis, reporting that “within-population genetic diversity is being lost (globally)” based on a “small but significant” effect size *g =* −0.11 (95% HPD *=* −0.15, −0.07).

Re-analyzing Shaw’s dataset, I show: (1) percentage genetic diversity change is not robust and close to zero, (2) most observations lack sufficient signal-to-noise ratio to detect diversity changes, and (3) evolutionary theory precisely predicts genetic diversity declines are expected to be small, but warns of invisibly growing large future losses.

## Understand genetic diversity loss as a percentage

Measuring global genetic diversity with interpretable values is paramount—especially as the UN CBD published a preliminary target to “protect 90% of genetic diversity within [every] species” ^8^, later removed due to lack of support ^8,10^. Shaw uses effect sizes with unclear biological meaning: effect size, *g* =(g*_*present*_ *-g*_*past*_*) /s*_*g*_ *× J*, is calculated with *g* being any genetic metric measured in two timepoints, *s*_*g*_ is the pooled standard deviation, and *J* is a sample size correction factor, and thus the effect size unit is the standard deviation of measurements of the different metrics. They further combine standard diversity metrics (see below) with other composition metrics like allele frequency, or population differentiation (*F*_*ST*_), or number of breeding adults (**Table S1, S2**). While such composition metrics are important essential variables describing different genetic or population processes, they are not measuring diversity, and increases or decreases may not even clearly be detrimental. The two key metrics of genetic diversity in a population are *average genetic diversity π* (equivalent to the species inverse Simpson diversity evenness) and *allelic richness, M* (equivalent to species richness). Previous studies that informed policy reported interpretable percentage losses of these diversity metrics: Leigh et al. ^5^ found ∼6% π and *M* decline over 97 years across 91 species, Millette et al. ^6^ measured no significant π decline in a mitochondrial gene over 36 years in 17,082 animal species, and Exposito-Alonso et al. ^8^ and Mualim et al. ^11^ inferred ∼10–20% π loss and ∼30–40% *M* loss using diversity-area relationships trained with genome-wide data of ∼30 species and projected to 13,808 species.

To translate Shaw’s effect size decline into a percentage diversity change, I first filtered observations to diversity metrics (*n*=916 π for 323 species, *n*=785 M for 283 species), removed coalescent studies (not direct temporal measures), subsetted to nuclear markers, removed domesticated species/pests, and excluded outliers (see steps and rationale in **Supplemental Materials**. NB: conclusions below are not driven by subsetting, rather provide a clearer signal, **Tables S3, S4**).

With this standardized diversity metric subset of the dataset, I quantified actual genetic diversity changes from past to present as a percentage (*%π* = 100*×* (*π*_*present*_ *-π* _*past*_)*/ π* _*past*_). I found no clear evidence of diversity change in any direction (**Fig. 1A-B, Fig. S2, S4, S5, S7, S8, S11, S12**). Three averaging methods yielded non-robust results near zero. The arithmetic average showed non-significant increase (%π = +0.99% ±1.94% [95% CI]), the study error-weighted average showed significant increase (+0.16% ± 0.09% [95% CI]), while meta-regression varied from a significant negative trend (−1.40% ± 1.02% [95% HPI]), to a non-significant trend (−0.78% ± 1.08% [95% HPI]), to a significantly positive trend (0.21% ± 0.10% [95% HPI]) (see other models, **Table S5**, similar results for *M* in **Table S8**, and geometric average of π and *M*, **Tables S7, S9**). Even conducting analyses using the Hedge’s g effect size from Shaw, we see the arithmetic average to be negative (*g =* −0.057 ± 0.038 [95% CI]), the study error-weighted average showed a significant increase (*g =* +0.01 ± 0.00011 [95% CI]), and their own meta-regression model shows a significant decline trend (*g =* −0.05 ± 0.03 [95% HPI]) becoming non-significant when changing priors or accounting for further background random effects such as species group and genetic marker type (see 32 model variations, **Table S2**). The reason I reached a different conclusion than Shaw is that studying the data on the scale of percent diversity revealed very small single digit percent average changes, which may be strongly affected by analysis method and data quality. Conclusions must thus uphold across multiple averaging methods, and much attention to data quality and null expectations is required (see below). My analyses demonstrate no robust statistical signal.

**Fig. 1.**
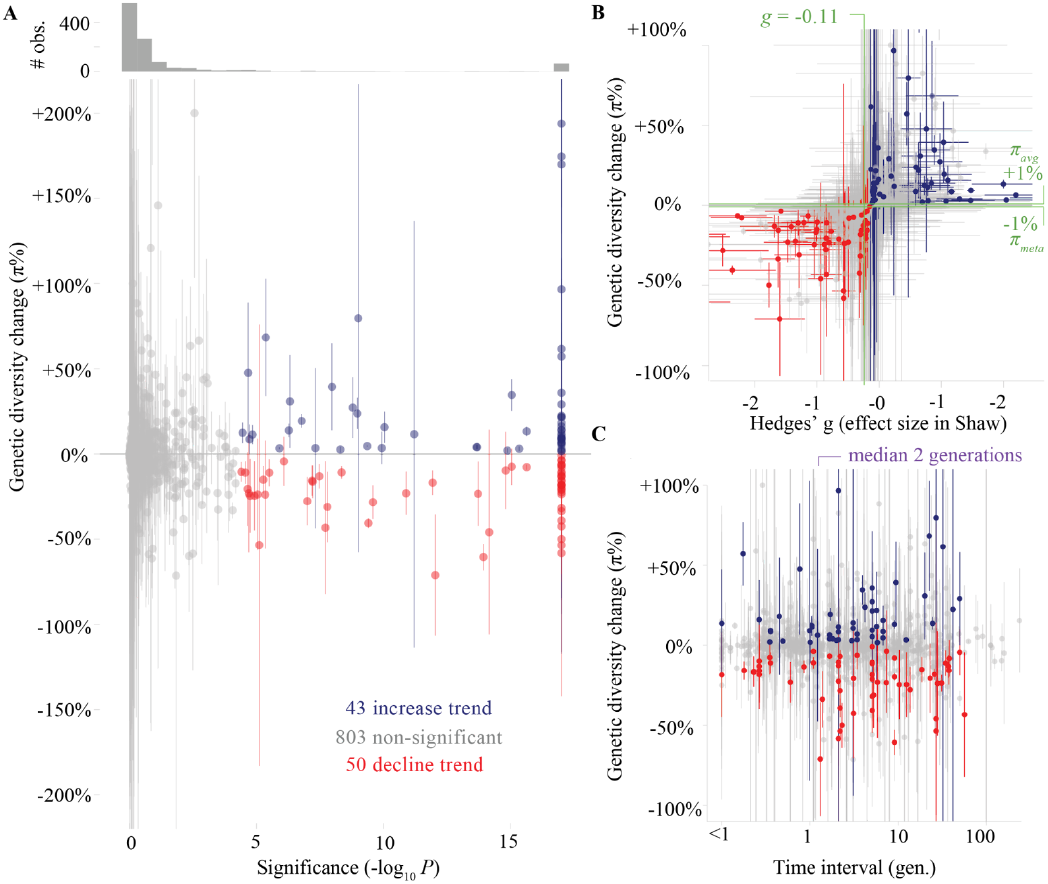
No robust evidence for global genetic diversity decline. **(a)** Raw genetic diversity change (π and He) between two time points (916 studies, 323 species) and the corresponding *P*-value of significance conducting per-study Hedge’s g using a t-distribution and family wise test correction (Bonferroni, *P ×* number of tests). **(b)** Comparison between original Hedge’s g in Shaw and actual raw genetic diversity change %, highlighting in green the average reported *g=* −0.11 by Shaw and several re-calculated %π averages (see main text). **(c)** Genetic diversity change %π across study time intervals.

To further understand why the average diversity trend is centered around zero, I analyzed studies separately. I found that there were roughly equal upward and downward trajectories across the entire dataset (**Text S2**, Binomial test could not reject 50/50% chance in all cases, *n*=4,023, 628 species, *P* = 0.6). Testing each diversity observation Hedges’ g trajectory using t test and Bonferroni correction (n= 916 %π, 323 species, **Fig. S3, Text S2, S3**), I found that only ∼10% of the studies in the Shaw π dataset produced significant results, with roughly ∼5% positive diversity trends and ∼5% negative diversity trends (*n*_positive_= 43, *n*_negative_ *=* 50, Binomial test *P*= 0.5. Identical results for *M* diversity, even when it is expected to be more sensitive to change ^8,12^).

### Understanding the signal-to-noise in genetic diversity datasets

Seemingly random upward and downward genetic diversity trends may reflect limited sampling rather than true population changes. Population genetic simulations of stable populations matching Shaw’s sampling efforts (**Text S4, Fig. S9, S10**) produced expected average diversity changes from sampling alone of π% = −0.81 to +2.4% across 1,000 bootstraps—similar range to observed trends (**Fig. S15, S16**). Simulations indicate that more than 50% reviewed in Shaw contain at least ±17%π measurement error (95%CI) (**Fig. S17**, note *n*_*median*_ *∼* 40 individuals, *L*_*median*_ *∼* 10 markers).

While sampling variation explains most trends, some species may show clear decline signals. Examining the extreme tail (5%) of the null distribution, I found 22 species with genetic diversity declines beyond sampling expectations (**Fig. 1C,D**), averaging %π= −18% decline compared to −8.7% from just sampling. Notable among these is *Strigops habroptila*, the critically endangered New Zealand parrot kākāpō, whose decline has been widely documented.

### Understanding why genetic diversity loss occurs slowly from evolutionary genetic theory

Despite the lack of robust consistent statistical trends, genetic diversity is likely declining in many species: documented demographic bottlenecks in conservation indices ^13^ must impact genetic diversity ^8^. However, the key questions are: *what is the expected genetic diversity loss across species given these demographic impacts?* Evolutionary genetics predicts small changes over short time intervals (median = 2 generations in Shaw’s dataset), which become substantial long-term ^11^.

Applying evolutionary theory in various population contraction scenarios, we can express genetic diversity as population size (*N*_*0*_ *→ N*_*1*_) or geographic range (*A*_*0*_ *→ A*_*1*_) losses (**Mathematical Appendix**).

A common conservation scenario concerns a local population bottleneck *N*_*0*_ *→ N*_*1*_, which follows: *π*_*t*_ *= π*_*0*_ *×(1-1/2N)*^*t*^, which rearranged yields: *N*_*1*_ *= 1 / (2 × (1 -(1-X*_*%*π_)^*1/t*^ )). This provides the intuition that to generate a −10%π diversity change we require a substantial bottleneck of ∼24 individuals for ∼5 generations—unlikely across most species. Species showing significant negative trends may have suffered bottlenecks of *N*_*1* median_ = 2,570 individuals accounting for species-specific generation intervals (**Fig. 2, Fig. S20, Text S4**). Some extreme cases like kākāpō are known to have experienced even more severe bottlenecks.

**Fig. 2.**
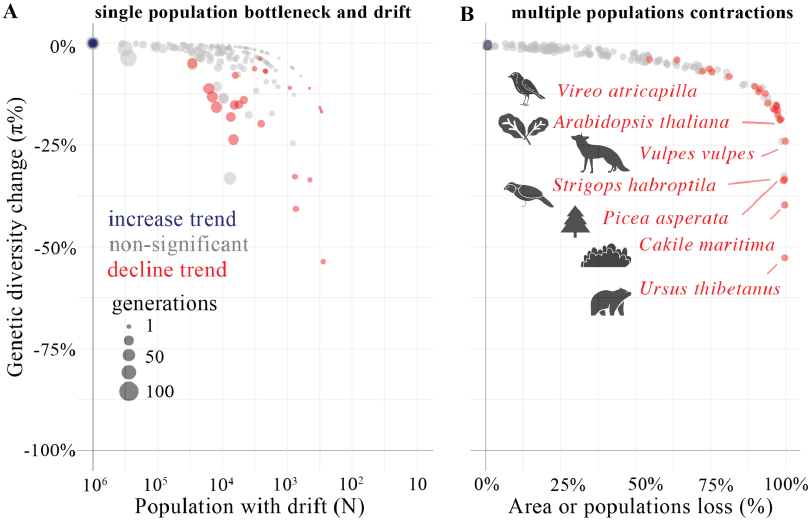
Extreme population crashes required to explain substantial % genetic changes. **(a)** Population bottleneck size required to explain π% change trends from genetic drift over generations. The number of generations assigned to each species temporal dataset from the original data is used for prediction. (**b**) Species geographic area or meta-population size loss required to explain π% change trends. This is independent of the starting population size, and occurs even if the remaining populations and overall species census are still large.

Alternatively, global habitat contractions eliminate entire populations with unique genetic diversity. Using the genetic-diversity-area relationships ^8,11^ (GDAR/MAR) that follows: *X*_*%*π_*=1-(1-A*_*1*_ */A*_*0*_)^*z*^; where *z ∼* 0.05 conservatively captures among-groups spatial genetic structure ^8^. We can re-arrange this to: *X*_*A*_ *= 1-A*_*1*_*/A*_*0*_ *= 1-(1-X*_*%*π_)^*1/z*^. This gives the intuition that a −10%π diversity change requires a 87% habitat area loss (**Fig. 2, Fig. S22, Text S4**). These extreme requirements confirm that large genetic diversity losses are not expected short-term unless complete geographic range crashes have occurred.

Detecting genetic diversity changes below 10% would require substantially larger sampling efforts than currently employed. For instance, simulations show sampling of ∼200 individuals/timepoint and 20,000 variants achieves ±1%π accuracy (95%CI) (**Fig. S17**).

Finally, Shaw studies which conservation actions increase genetic diversity. Roughly 50% of observations showed increasing diversity trends. Genetic diversity does not increase from natural population recovery in the time scales represented in the Shaw dataset—diversity grows slower than it decreases (**Fig. S26, Text S5**). Positive trends require unrealistic population growth of ∼60 fold (**Fig. S25**). Instead, artificial mixing of differentiated populations (*F*_*ST*_) explains increases (**Fig. S16, S27**). Shaw’s finding that “supplementation” (i.e. adding individuals to an existing population by translocation) was the only conservation factor significantly associated with positive trends supports this. While supplementation may be necessary sometimes ^3^, population mixing cannot be prescribed generally and can even harm threatened populations ^14^. Positive trends may therefore mask, rather than indicate, conservation success.

## Conclusion

Our inability to measure genetic diversity loss across all species offers a silver lining: there is still time to protect genetic diversity. However, the lack of robust signal in my re-analyses does not mean genetic diversity is safe. This meta-analysis mirrors counterintuitive and complex trends in other biodiversity meta-analyses. While extinctions are increasing, studies of species richness in century-long meta-analyses showed no systematic loss of local species numbers, with equal positive and negative trends across ecosystems ^15,16^. Population decline debates follow similar patterns: the Living Planet Index shows half of censused populations do not change or even increase ^18^. Yet extinctions and population collapses are undeniable ^13,19^. Genetic diversity is likely declining globally, but evolutionary theory predicts only small percentage losses at short timescales. Many current public datasets lack sufficient signal-to-noise ratios to detect such changes due to small sample sizes and obsolete DNA technologies. However, current non-significant or weak genetic losses are temporary—evolutionary theory predicts genetic diversity will continue declining for hundreds to thousands of generations until roughly matching current population losses ^11^. We face an enormous genetic diversity extinction debt, and to prevent irreversible and measurable genetic diversity losses we must protect and recover populations now.

## Data and code availability

All data and code to replicate these analyses are available at: github.com/moiexpositoalonsolab/gendivmeta.

## Acknowledgements

I am deeply thankful to Shaw et al. for their enormous efforts to compile genetic data across the literature, and for their thoughtful analyses. In addition, I thank Shaw and Grueber for their feedback while corresponding about these analyses. I am grateful for feedback on the manuscript from Jarrod Hadfield, John Huelsenbeck, Patricia Lang, Meixi Lin, Katie Millette, Rasmus Nielsen, David Reich, Miles Roberts, Peter Sudmant, Jeff Spence, Perry Valpine, and Detlef Weigel, and for discussions with members of the MOILAB.

## Supplemental Materials

**Tex S1-S7**

**Fig. S1-28**

**Tables S1-8**

**Response-GlobalMetaAnalysisGenetics.pdf**

## Supplemental Mathematical Appendix

**Loss scenario 1a, 1b, 2a**,**2b**

**Gain scenario 1b, 2a**

**Response-Mathematical-Appendix.pdf**

## Supplemental Materials | Response to Shaw et al. (2025)

### Text S1: Subset dataset to robust and comparable genetic diversity metrics (π and S)

#### Summary of numbers from the original dataset

Shaw et al (2025) include a number of metrics of genetic composition in their analyses. The definition (from Authors) is below:

AR (allelic richness) A (mean alleles)

TA (total alleles)

pA (total private alleles)

NPL (number of polymorphic loci)

Other1 (other metric associated with variant counts)

He (expected heterozygosity)

pi (nucleotide diversity) h (haplotype diversity)

H (Shannon diversity index)

PIC (polymorphic information content) NEA (number of effective alleles)

Fis (inbreeding coefficient)

R (mean relatedness/kinship among individuals)

apFst (among-population fixation index)

Freq (frequency of an allele of interest)

NPD (mean individual nucleotide p-distance)

BS (bandsharing score)

Other2 (other metric associated with Variant frequencies)

Ho (observed heterozygosity)

SH (standardised multilocus heterozygosity)

Ai (mean alleles per individual)

F (inbreeding coefficient at the individual level)

NH (number of polymorphic loci per individual)

Other3 (other metric associated with Individual-level diversity)

Ne (effective population size estimated from molecular data)

Nd (effective population size estimated from demographic data)

Nb (effective number of breeders),

Nf (female effective population size),

Nc (effective population size estimated from a population census, and calculated based on an assumption about the Ne:Nc ratio),

Other4 (other metric associated with integrated statistics).

These were further grouped in four categories (definitions from Authors):

Group1 = metrics based on variant counts (AR, A, TA, pA, NPL, Other1)

Group2 = metrics based on evenness of variant frequencies (He, pi, h, H, PIC, NEA, Fis, R, apFst, Freq, NPD, BS, Other2)

Group3 = metrics based on population means of individual-level diversity (Ho, SH, Ai, F, NH, Other3)

Group4 = integrated statistics (Ne, Nd, Nb, Nf, Nc, Other4).

In this reanalyses, the dataset is filtered to only standard metrics of diversity:

**Table S 1:**
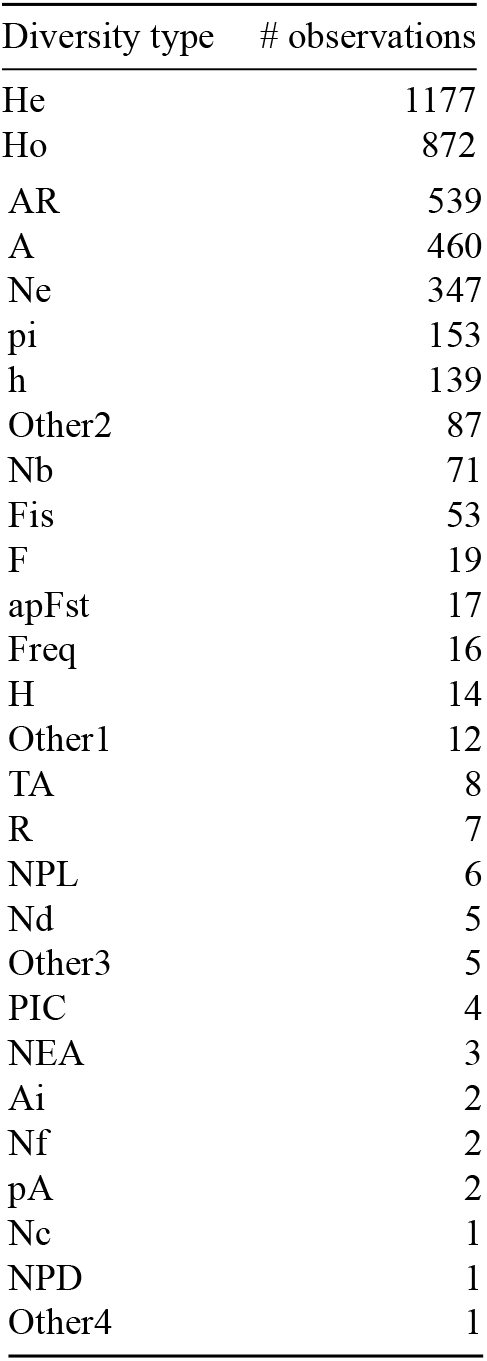
Number of observations by diversity type.

- We then focus on the one hand on average difference metrics, π diversity, He and π diversity. There are a total of 1177 and 153 studies with He and Pi measurements. We decided to combine them so there is sufficient sample size for downstream analyses (for biallelic loci, expected heterozygosity and π are the same value). We also focus on the number of alleles, 999, for which we also have population genetic expectations. Together, these are 57.8921203% of original dataset.
- We also focus on studies with metrics in 2 time points, rather than linear trajectories or coalescent, which makes comparisons most straightforward. The number of 2 timepoint measurements is: 3803, linear: 64, and coalescent: 156. Coalescent is 3.8777032% of the original dataset.
- The number of observations by Kingdom and Phylum mostly show observations include insects, vertebrates, and flowering plants. We do not do any filtering.
- Number of observations of domesticated species: Domesticated 767 observations (% 19.0653741). Domesticated/pathogens/pests 3256, 767 observations (% 80.9346259, 19.0653741). We removed domesticated species.
- Number of nuclear vs other genomes: Nuclear genome markers 3658 observations (% 90.9271688). We focus on nuclear, as chloroplast and mitochondria undergo different evolutionary dynamics (maternal inheritanze, no recombination during meiosis). There are 3658 nuclear, 282 mitochondrial, and 13 mixed genomes. Non-nuclear is 9.0728312% of original dataset.

#### Summary of numbers from filtered dataset

Total number 1701 of which 54% are π. The final subset is 42% of the original dataset.

Summaries of allelic richness π

- 916 studies
- 323 species
- Summary of number of generations in the final dataset is: Mean=6.5 generations, Median=2 generations, IQR= 0.59
- 6 generations.

Summaries of allelic richness *M*

- 785 studies
- 283 species
- Summary of number of generations in the final dataset is: Mean=6.8 generations, Median=2 generations, IQR= 0.62
- 6.9 generations.

**Figure S 1:**
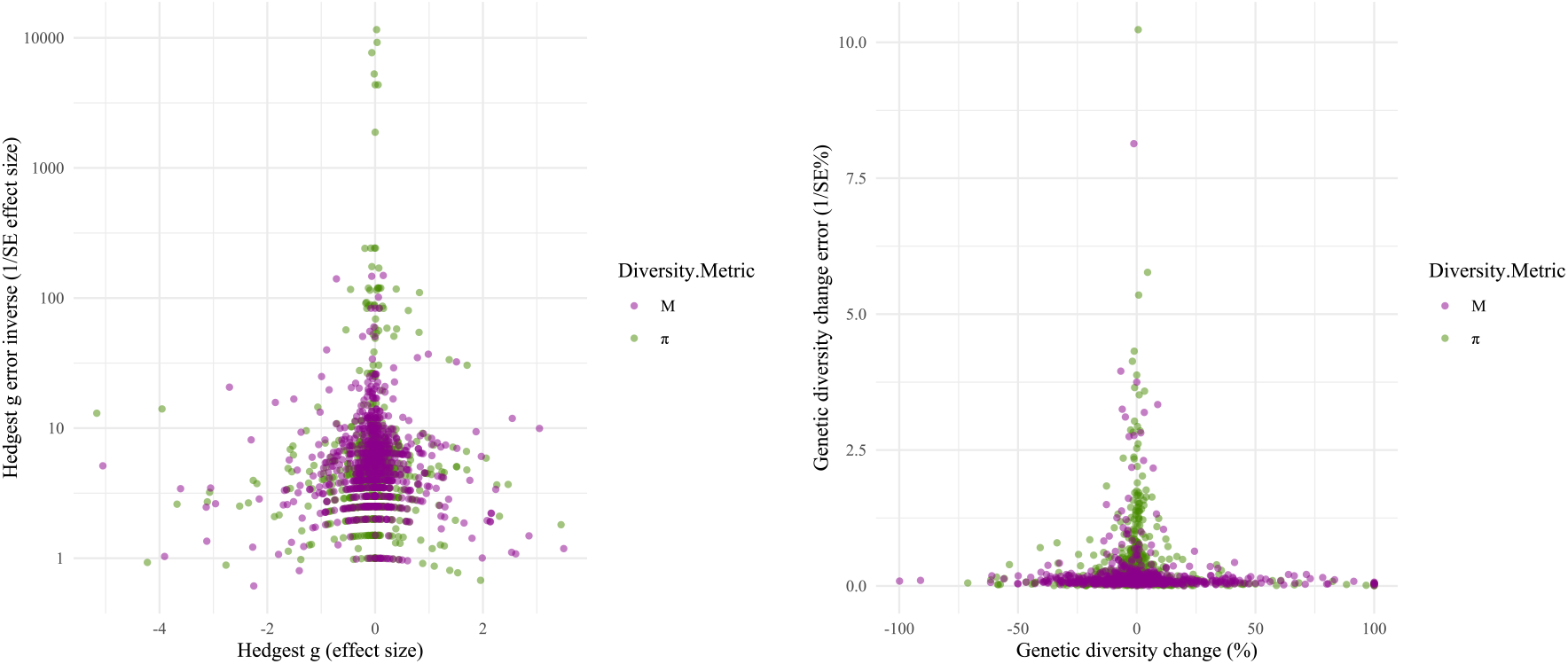
Funnel plot of effect size vs error shows a classic relationship of smaller effect size in measurements with less error.

Figure S 1 shows the typical funnel shape whereby most precise estimates (high 1/SE) show smaller effects.

### Re-calculate effect sizes

To reproduce Shaw calculation of effect size:

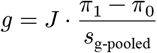

To calculate the corrected version we have the degrees of freedom used for effect size correction is: *df* = *n*_early_ + *n*_recent_ − 2. We used Hedges’ correction factor *J* for small-sample bias: 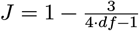.

Its standard error was estimated as:

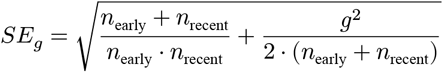

### Calculate a percentage diversity change

#### Percentage change

The fraction (or percentage if multiplied × 100) of diversity change is defined as:

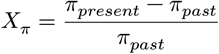

Each estimate came with sample size and some form of error extracted during the review process for both past (early) and present (recent) timepoints. The errors reported came in several forms: standard deviation (SD), standard error (SE), or the width of a 95% confidence interval (CI). All error types were converted to standard errors using the following rules (diversity π and *M* had mostly errors reported as SD or SE, only a few 95%CI and only 6 observations of *M* with IQR, discarded):

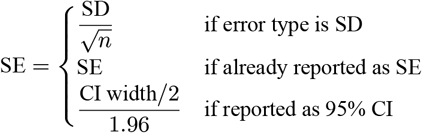

where *n* is the sample size. From this standard deviation (SD) was recomputed for completeness, as sample sizes were known for all observations as 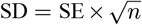. This was done separately for early and recent time points. Then pooled standard deviation was computed as:

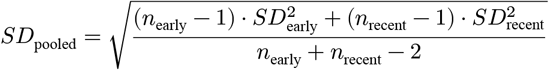

Just to have something informative of both studies, the effective sample size was computed as the harmonic mean of early and recent sample sizes: 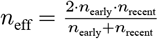.

From this, we aim to compute a error of the measured change 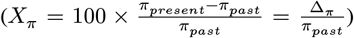 The standard error of the change is computed assuming errors in the two time points are independent and estimated the standard error of the difference: 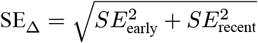, and scaled to be percentage change error: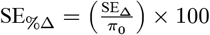.

This standard error is not exact because of the division of π_0_, which itself contains noise. We can attempt to create a second standard error uses the delta method. While both are approximations, the delta method attempts to provide a more sophisticated approximation. Consider 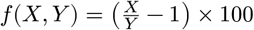, where *X* = π and *Y* = π_0_. We apply the **delta method** to approximate the variance of *f*(*X, Y*). The first-order Taylor expansion is:

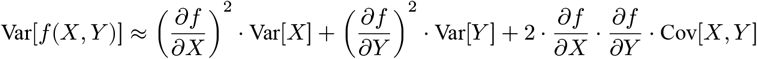

Assuming independence between *X* and *Y* (i.e., Cov[*X, Y*] = 0), the partial derivatives are:

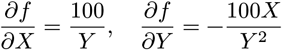

Substituting into the variance expression:

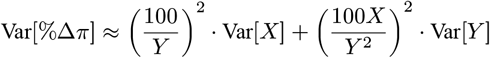

Thus, the **standard error** of the percent change is:

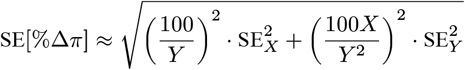

#### Log ratio change

Although likely an arithmetic percentage change is the most understandable metric, for completeness, I also calculate the ratio of change:

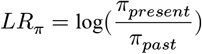

When averaging the log ratio we get a geometric (rather than arithmetic) mean, which may be helpful.

We can also calculate the standard error of the log ratio using the delta method. Let: *X* = π (recent diversity) and *Y* = π_0_ (early diversity), and 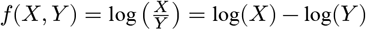. We aim to compute the standard error of the log-ratio, 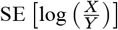, using the **delta method**, assuming that *X* and *Y* are independent.

The variance of *f*(*X, Y*) is approximated as:

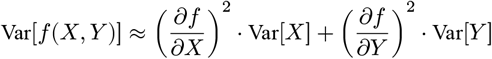

The partial derivatives are:

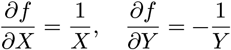

Substituting into the variance approximation:

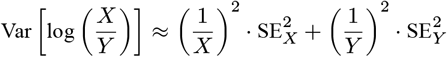

Therefore, the standard error of the log-ratio is:

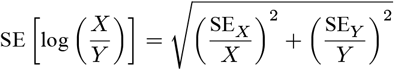

### Summary statistics of changes

Summaries of π change:

Median [IQR] Hedges g = −0.026 [-0.17 - 0.1]

Median [IQR] Error of Hedges g = 0.22 [0.15 - 0.33]

Median [IQR] Percentage change = −0.78% [-5 - 2.8%]

Median [IQR] SE of Percentage change = 7.4% [3.5 - 16%]

Mean Percentage change= 0.99%

Mean SE of Percentage change= 14%

Summaries of *M* change:

Median [IQR] Hedges g = −0.0071 [-0.23 - 0.16]

Median [IQR] Error of Hedges g = 0.2 [0.13 - 0.3]

Median [IQR] Percentage change = −0.44% [-7.9 - 6.8%]

Median [IQR] SE of Percentage change = 10% [6.5 - 16%]

Mean Percentage change= 1.5%

**Figure S 2:**
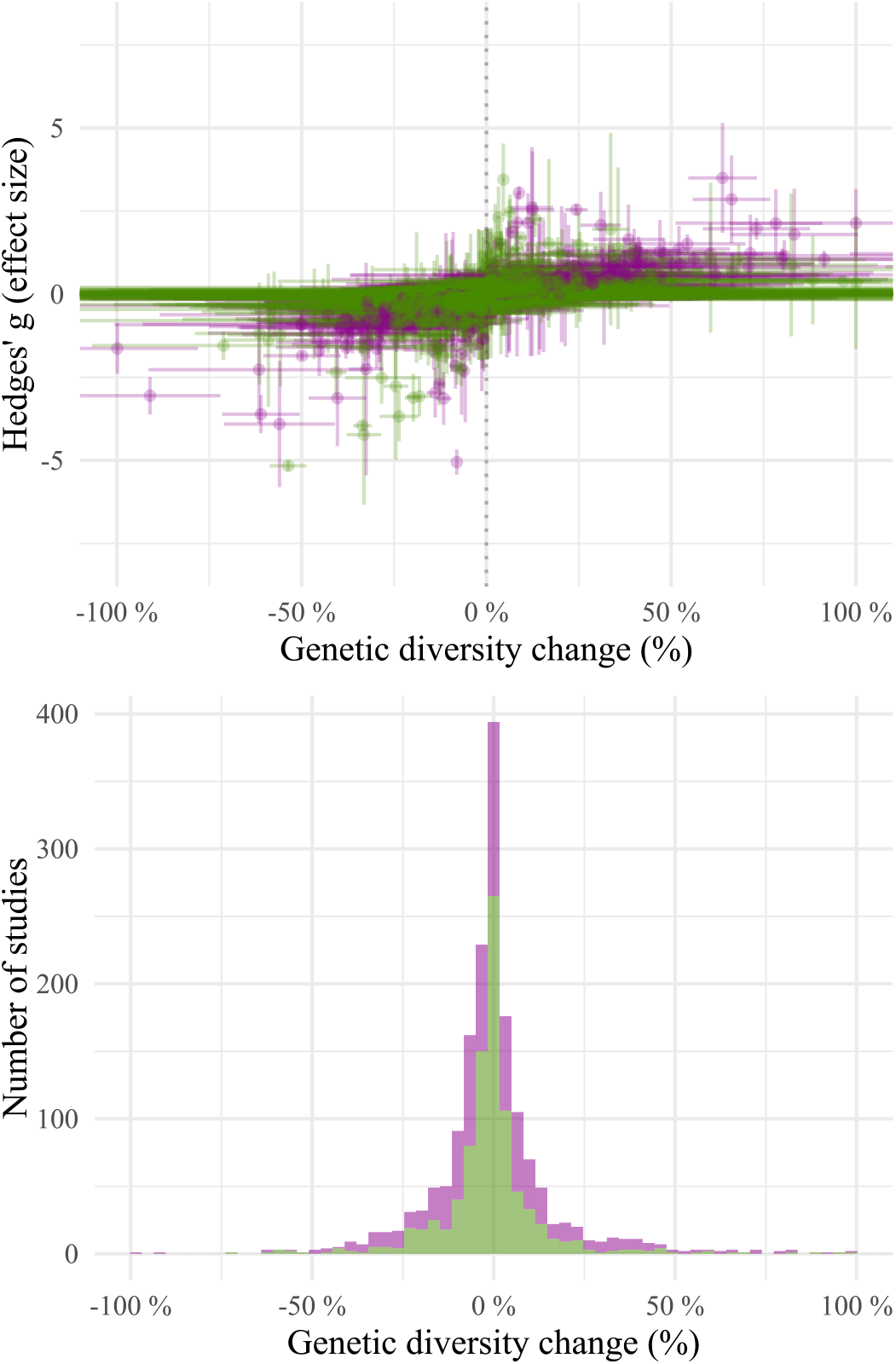
Distribution of effect sizes (g) and raw genetic diversity changes (%).

Hedges’ g and genetic diversity change in percentage are correlated since raw data is used to calculate g (Figure S2), although not perfectly (Pearson’s r=0.53). The extremest values in π, which comprises the majority of the data however, typically do not coincide. We can see that of the 5% top π values that decline based on Hedge’s g (46), only a small fraction are also among the top raw % values of decline (43%). Same for *M* (50%). Figure S5 shows the raw average and estimated effect sizes in the comparision of Hedges’ g and raw values of change (colors indicate significance, see section below).

### Text S2: Test the temporal shift of genetic diversity changes

#### Calculate average change with different methods

*For diversity* π

Arithmetic mean Hedge’s g = −0.057 ±0.038 (95% CI).

Weighted SE mean Hedge’s g = 0.01 ± 0.00011(95% CI) (z = 194, *P* = 0).

MCMCglmm mean (intercept) of Hedge’s g = −0.05 ± 0.03 (95% HPI).

Arithmetic mean % π = 0.99% ± 1.9% (95% CI).

Weighted SE mean % π = 0.16% ± 0.089% (95% CI) (z = 3.6, *P* = 3.3515656 × 10^−4^).

MCMCglmm mean (intercept) of % π = −1.20 ± 0.73 (95% HPI).

Geometric mean % π = −1.2% ± 1.2% (95% CI).

Weighted SE geometric mean % π = 0.25% ± 0.089% (95% CI).

*For allelic richness M*

Average Hedge’s g = −0.044 ±0.049 (95% CI).

Weighted SE mean Hedges’g = −0.087 ± 0.005(95% CI) (z = −34, *P* = 4.3656128 × 10^−258^).

MCMCglmm mean (intercept) of Hedges’g = −0.04 ± 0.04 (95% HPI).

Arithmetic mean % *M* = 1.5% ± 1.9% (95% CI).

Weighted SE mean % *M* = −1.1 ± 0.13(95% CI) (z = −16, *P* = 1.0272574 × 10^−55^).

MCMCglmm mean (intercept) of % *M* = 0.46 ± 1.57 (95% HPI).

Geometric mean % π = −1.8% ± 2.3% (95% CI).

Weighted SE geometric mean % π = −0.86% ± 0.13% (95% CI).

#### Variations of meta-regression

To study the robustness of the original MCMCglmm meta-regression from Shaw, we conduct several variations of their function. From their supplement, they run MCMCglmm(g ∼ Z.year.Midpoint + Z.Gen.interval, random= ∼PaperID) using weak priors. Following Shaw, I used a weak inverse-gamma model prior (V=1, nu=0.002) Aka. “pweak”.

I generated variations over this model to understand robustness and explain why the results may differ from an arith-metic or error-weigthed mean (models were named numerically with increasing fixed or random terms, “m1”…”m6”):

Changing fixed effects. I first changed the mean-centered generation interval fixed effect (shaw1 model). The mean-centering is necessary so that the intercept corresponds with the average fixed effect variable. However, the generation interval was not normally distributed. In fact, >75% of the dataset showed lower number of generations (quantile 75% = 6 generations) than the mean (6.5 generations). This is due to a long tail effect. A approach less driven by long tails is including a fixed effect of generation interval median-centered. When conducting that, the effect typically becomes much weaker or non-significant (this corrected model was named shaw2).

Changing random effects. Shaw includes Paper ID, and I tested the inclusion of Kingdom, species, and Marker type, to account for other possible heterogeneity in the data.

Changing priors. Although a weak prior seems sensible, in Bayesian statistics is important to test model robustness based on priors, especially when signals appear weak. I tested not including any prior (using default MCMCglmm, nu=0, V=1, aka. “none”). I tested a more regularizing prior “preg” (V=1e-4, nu=10), and a prior that would limit unexplained variance to focus on the observation measurement error term (1/*SE*^2^), aka “pfix” (V=1e-4, nu=100).

The last one is closer to fixed effect meta-analysis. These priors represent a range from low to strong shrinking, from weak information but unstable estimates to stronger information in model from standard error measurements.

The same set of models was fitted to the entire dataset (using effect sizes) as well as standard average diversity π and allelic richness *M* both effect sizes and percentage change. Models in reporting tables are ranked by DIC from MCMCglmm.

Several top models in both π and *M* percentage changes included non-significant, positive and negatively significant trends.

#### All Shaw data effect size

**Table S 2:** Shaw effect size meta-regression model comparison.

| label | DIC | summary | fixed effects | random effects |
| --- | --- | --- | --- | --- |
| m5_none | 8874.6 | -0.03 ± 0.28 ns | M.Gen.interval | PaperID + Kingdom + Binomen.rotl |
| m5_pweak | 8873.8 | -0.03 ± 0.27 ns | M.Gen.interval | PaperID + Kingdom + Binomen.rotl |
| mshaw2_pfix | 10080.5 | -0.05 ± 0.03 * | M.Year.Midpoint + M.Gen.interval | PaperID |
| m5_pfix | 10087.2 | -0.05 ± 0.03 * | M.Gen.interval | PaperID + Kingdom + Binomen.rotl |
| m3_preg | 10110.6 | -0.06 ± 0.03 * | M.Gen.interval | none |
| m3_pfix | 10087.0 | -0.06 ± 0.03 * | M.Gen.interval | none |
| m4_pfix | 10087.5 | -0.06 ± 0.03 * | M.Gen.interval | PaperID |
| m3_none | 10112.9 | -0.06 ± 0.03 * | M.Gen.interval | none |
| m3_pweak | 10113.3 | -0.06 ± 0.03 * | M.Gen.interval | none |
| m2_pfix | 10086.5 | -0.06 ± 0.03 * | none | PaperID |
| m6_pfix | 10086.2 | -0.06 ± 0.03 * | M.Gen.interval | PaperID + Kingdom + Binomen.rotl + Marker.Type |
| m1_pweak | 10112.1 | -0.06 ± 0.03 * | none | none |
| mshaw1_pfix | 10080.6 | -0.06 ± 0.03 * | Z.Year.Midpoint + Z.Gen.interval | PaperID |
| m1_pfix | 10087.1 | -0.06 ± 0.03 * | none | none |
| m1_preg | 10109.5 | -0.06 ± 0.03 * | none | none |
| m1_none | 10112.6 | -0.06 ± 0.03 * | none | none |
| mshaw2_preg | 8877.4 | -0.11 ± 0.05 * | M.Year.Midpoint + M.Gen.interval | PaperID |
| mshaw2_pweak | 8878.5 | -0.11 ± 0.05 * | M.Year.Midpoint + M.Gen.interval | PaperID |
| mshaw2_none | 8878.0 | -0.11 ± 0.05 * | M.Year.Midpoint + M.Gen.interval | PaperID |
| mshaw1_preg | 8876.9 | -0.11 ± 0.05 * | Z.Year.Midpoint + Z.Gen.interval | PaperID |
| mshaw1_pweak | 8878.4 | -0.11 ± 0.05 * | Z.Year.Midpoint + Z.Gen.interval | PaperID |
| mshaw1_none | 8878.4 | -0.11 ± 0.05 * | Z.Year.Midpoint + Z.Gen.interval | PaperID |
| m5_preg | 8872.7 | -0.11 ± 0.05 * | M.Gen.interval | PaperID + Kingdom + Binomen.rotl |
| m4_preg | 8873.2 | -0.12 ± 0.05 * | M.Gen.interval | PaperID |
| m4_none | 8874.2 | -0.12 ± 0.05 * | M.Gen.interval | PaperID |
| m4_pweak | 8874.3 | -0.12 ± 0.05 * | M.Gen.interval | PaperID |
| m2_preg | 8871.8 | -0.12 ± 0.05 * | none | PaperID |
| m2_none | 8873.9 | -0.12 ± 0.05 * | none | PaperID |
| m2_pweak | 8873.5 | -0.12 ± 0.05 * | none | PaperID |
| m6_none | 8782.2 | -0.25 ± 0.46 ns | M.Gen.interval | PaperID + Kingdom + Binomen.rotl + Marker.Type |
| m6_pweak | 8780.4 | -0.25 ± 0.47 ns | M.Gen.interval | PaperID + Kingdom + Binomen.rotl + Marker.Type |
| m6_preg | 8804.7 | -0.30 ± 0.18 * | M.Gen.interval | PaperID + Kingdom + Binomen.rotl + Marker.Type |

The exact model from Shaw et al. (“mshaw_pweak”) recapitulates the found g=-0.11 reported int he paper. We note that the fixed erffect generation interval is non-significant, indicating that with increasing generation interval there is a positive trend in the effect size.

**Table S 3:** Shaw meta-regression summary.

|  | post.mean | l-95% CI | u-95% CI | eff.samp | pMCMC |
| --- | --- | --- | --- | --- | --- |
| (Intercept) | -0.1121301 | -0.1649134 | -0.0621774 | 1166.092 | 0.001 |
| Z.Gen.interval | 0.0344938 | -0.0388307 | 0.1004779 | 1000.000 | 0.346 |
| Z.Year.Midpoint | 0.0948754 | 0.0216149 | 0.1693552 | 1000.000 | 0.018 |

Exclusion of the key diversity variables makes the Shaw model non-significant.

**Table S 4:** Shaw meta-regression summary excluding π and M.

|  | post.mean | l-95% CI | u-95% CI | eff.samp | pMCMC |
| --- | --- | --- | --- | --- | --- |
| (Intercept) | -0.0578699 | -0.1316435 | 0.0082411 | 1000 | 0.106 |
| Z.Gen.interval | 0.0357412 | -0.0396524 | 0.1116065 | 1000 | 0.364 |
| Z.Year.Midpoint | 0.1116704 | 0.0394472 | 0.2072596 | 1000 | 0.012 |

#### *Average genetic diversity* π percent change

**Table S 5:** Percentage diversity π meta-regression model comparison.

| label | DIC | summary | fixed effects | random effects |
| --- | --- | --- | --- | --- |
| mshaw1_pfix | -4852.1 | $0.26 \pm 0.32$ ns | Z.Year.Midpoint + Z.Gen.interval | PaperID |
| mshaw2_preg | -4856.6 | $0.24 \pm 1.24$ ns | M.Year.Midpoint + M.Gen.interval | PaperID |
| mshaw2_pfix | -4852.0 | $0.21 \pm 0.10$ * | M.Year.Midpoint + M.Gen.interval | PaperID |
| m2_pfix | -4853.1 | $0.16 \pm 0.09$ * | none | PaperID |
| m1_pfix | -4852.7 | $0.16 \pm 0.09$ * | none | none |
| m3_pfix | -4853.6 | $0.13 \pm 0.09$ * | M.Gen.interval | none |
| m4_pfix | -4851.1 | $0.13 \pm 0.09$ * | M.Gen.interval | PaperID |
| m5_pfix | -4854.0 | $0.13 \pm 0.09$ * | M.Gen.interval | PaperID + Kingdom + Binomen.rotl |
| m6_pfix | -4852.2 | $0.13 \pm 0.09$ * | M.Gen.interval | PaperID + Kingdom + Binomen.rotl + Marker.Type |
| mshaw2_none | 6371.4 | $-0.78 \pm 1.08$ ns | M.Year.Midpoint + M.Gen.interval | PaperID |
| mshaw2_pweak | 6371.6 | $-0.78 \pm 1.11$ ns | M.Year.Midpoint + M.Gen.interval | PaperID |
| m6_preg | 6778.0 | $-1.05 \pm 0.72$ * | M.Gen.interval | PaperID + Kingdom + Binomen.rotl + Marker.Type |
| m3_pweak | 6794.5 | $-1.05 \pm 0.74$ * | M.Gen.interval | none |
| m3_none | 6795.0 | $-1.05 \pm 0.74$ * | M.Gen.interval | none |
| m4_preg | 6778.3 | $-1.05 \pm 0.74$ * | M.Gen.interval | PaperID |
| m3_preg | 6778.6 | $-1.06 \pm 0.72$ * | M.Gen.interval | none |
| m5_preg | 6010.0 | $-1.11 \pm 0.91$ * | M.Gen.interval | PaperID + Kingdom + Binomen.rotl |
| m1_none | 6797.9 | $-1.19 \pm 0.72$ * | none | none |
| m1_pweak | 6798.8 | $-1.19 \pm 0.72$ * | none | none |
| m2_preg | 6781.8 | $-1.19 \pm 0.71$ * | none | PaperID |
| m1_preg | 6782.3 | $-1.20 \pm 0.71$ * | none | none |
| m4_none | 6461.3 | $-1.27 \pm 1.06$ * | M.Gen.interval | PaperID |

Table S 5: Percentage diversity $\pi$ meta-regression model comparison. (continued)
| label | DIC | summary | fixed effects | random effects |
| --- | --- | --- | --- | --- |
| m4_pweak | 6462.4 | $-1.28 \pm 1.08$ * | M.Gen.interval | PaperID |
| m2_pweak | 6460.5 | $-1.40 \pm 1.03$ * | none | PaperID |
| m2_none | 6460.4 | $-1.40 \pm 1.02$ * | none | PaperID |
| m5_pweak | 6460.8 | $-1.93 \pm 4.75$ ns | M.Gen.interval | PaperID + Kingdom + Binomen.rotl |
| m5_none | 6462.3 | $-1.95 \pm 4.53$ ns | M.Gen.interval | PaperID + Kingdom + Binomen.rotl |
| mshaw1_preg | 6765.2 | $-3.12 \pm 1.52$ * | Z.Year.Midpoint + Z.Gen.interval | PaperID |
| mshaw1_none | 6371.1 | $-4.23 \pm 1.95$ * | Z.Year.Midpoint + Z.Gen.interval | PaperID |
| mshaw1_pweak | 6371.4 | $-4.23 \pm 1.96$ * | Z.Year.Midpoint + Z.Gen.interval | PaperID |
| m6_none | 6422.7 | $-4.47 \pm 8.30$ ns | M.Gen.interval | PaperID + Kingdom + Binomen.rotl<br>+ Marker.Type |
| m6_pweak | 6423.2 | $-4.54 \pm 8.45$ ns | M.Gen.interval | PaperID + Kingdom + Binomen.rotl<br>+ Marker.Type |

#### *Average genetic diversity* π percent change (delta method for SE)

**Table S 6:** Percentage diversity π (delta SE) meta-regression model comparison.

| label | DIC | summary | fixed effects | random effects |
| --- | --- | --- | --- | --- |
| mshaw2_pfix | -4848.3 | $0.05 \pm 0.10$ ns | M.Year.Midpoint + M.Gen.interval | PaperID |
| m2_pfix | -4835.7 | $-0.06 \pm 0.09$ ns | none | PaperID |
| m1_pfix | -4868.3 | $-0.06 \pm 0.09$ ns | none | none |
| m3_pfix | -4832.5 | $-0.06 \pm 0.09$ ns | M.Gen.interval | none |
| m6_pfix | -4838.4 | $-0.06 \pm 0.10$ ns | M.Gen.interval | PaperID + Kingdom + Binomen.rotl<br>+ Marker.Type |
| m5_pfix | -4847.1 | $-0.06 \pm 0.09$ ns | M.Gen.interval | PaperID + Kingdom + Binomen.rotl |
| m4_pfix | -4850.0 | $-0.06 \pm 0.09$ ns | M.Gen.interval | PaperID |
| mshaw1_pfix | -4836.6 | $-0.44 \pm 0.33$ * | Z.Year.Midpoint + Z.Gen.interval | PaperID |
| mshaw2_preg | -4807.6 | $-0.52 \pm 1.12$ ns | M.Year.Midpoint + M.Gen.interval | PaperID |
| mshaw2_pweak | 6376.6 | $-1.47 \pm 0.96$ * | M.Year.Midpoint + M.Gen.interval | PaperID |
| mshaw2_none | 6371.0 | $-1.48 \pm 0.97$ * | M.Year.Midpoint + M.Gen.interval | PaperID |
| m4_preg | 6777.3 | $-1.60 \pm 0.70$ * | M.Gen.interval | PaperID |
| m3_none | 6791.4 | $-1.60 \pm 0.72$ * | M.Gen.interval | none |
| m3_pweak | 6790.6 | $-1.62 \pm 0.69$ * | M.Gen.interval | none |
| m3_preg | 6777.8 | $-1.64 \pm 0.73$ * | M.Gen.interval | none |
| m1_pweak | 6795.3 | $-1.80 \pm 0.71$ * | none | none |
| m1_none | 6795.8 | $-1.82 \pm 0.71$ * | none | none |
| m1_preg | 6785.0 | $-1.82 \pm 0.69$ * | none | none |
| m4_pweak | 6456.3 | $-1.94 \pm 0.93$ * | M.Gen.interval | PaperID |
| m4_none | 6451.8 | $-1.94 \pm 1.03$ * | M.Gen.interval | PaperID |
| m2_pweak | 6453.0 | $-2.07 \pm 0.91$ * | none | PaperID |
| m2_preg | 6443.3 | $-2.11 \pm 0.90$ * | none | PaperID |
| m2_none | 6457.3 | $-2.13 \pm 0.90$ * | none | PaperID |
| m5_pweak | 6453.2 | $-2.70 \pm 5.61$ ns | M.Gen.interval | PaperID + Kingdom + Binomen.rotl |
| m5_preg | -4812.2 | $-3.01 \pm 1.06$ * | M.Gen.interval | PaperID + Kingdom + Binomen.rotl |
| m5_none | 6450.5 | $-3.03 \pm 5.67$ ns | M.Gen.interval | PaperID + Kingdom + Binomen.rotl |
| mshaw1_none | 6376.6 | $-4.93 \pm 1.75$ * | Z.Year.Midpoint + Z.Gen.interval | PaperID |

Table S 6: Percentage diversity $\pi$ (delta SE) meta-regression model comparison. (continued)
| label | DIC | summary | fixed effects | random effects |
| --- | --- | --- | --- | --- |
| mshaw1_pweak | 6369.6 | $-4.97 \pm 1.75$ * | Z.Year.Midpoint + Z.Gen.interval | PaperID |
| m6_pweak | 6413.6 | $-5.65 \pm 8.00$ ns | M.Gen.interval | PaperID + Kingdom + Binomen.rotl<br>+ Marker.Type |
| m6_none | 6409.2 | $-5.85 \pm 8.15$ ns | M.Gen.interval | PaperID + Kingdom + Binomen.rotl<br>+ Marker.Type |
| m6_preg | -4893.2 | $-6.24 \pm 6.16$ ns | M.Gen.interval | PaperID + Kingdom + Binomen.rotl<br>+ Marker.Type |
| mshaw1_preg | -4862.0 | $-10.04 \pm 1.61$ * | Z.Year.Midpoint + Z.Gen.interval | PaperID |

#### *Average genetic diversity* π log ratio (geometric)

**Table S 7:** Log ratio diversity π meta-regression model comparison.

| label | DIC | summary | fixed effects | random effects |
| --- | --- | --- | --- | --- |
| mshaw2_pfix | -1631.3 | $-0.01 \pm 0.01$ * | M.Year.Midpoint + M.Gen.interval | PaperID |
| mshaw2_pweak | -1762.9 | $-0.01 \pm 0.01$ ns | M.Year.Midpoint + M.Gen.interval | PaperID |
| mshaw2_none | -1767.6 | $-0.01 \pm 0.01$ ns | M.Year.Midpoint + M.Gen.interval | PaperID |
| mshaw2_preg | -1758.6 | $-0.01 \pm 0.01$ ns | M.Year.Midpoint + M.Gen.interval | PaperID |
| m3_none | -1489.4 | $-0.01 \pm 0.01$ * | M.Gen.interval | none |
| m4_pfix | -1615.2 | $-0.01 \pm 0.01$ * | M.Gen.interval | PaperID |
| m3_pfix | -1611.3 | $-0.01 \pm 0.01$ * | M.Gen.interval | none |
| m3_pweak | -1489.9 | $-0.01 \pm 0.01$ * | M.Gen.interval | none |
| m3_preg | -1505.4 | $-0.01 \pm 0.01$ * | M.Gen.interval | none |
| m4_preg | -1665.7 | $-0.01 \pm 0.01$ * | M.Gen.interval | PaperID |
| m2_pfix | -1610.3 | $-0.01 \pm 0.01$ * | none | PaperID |
| m1_pfix | -1607.3 | $-0.01 \pm 0.01$ * | none | none |
| m1_pweak | -1486.1 | $-0.01 \pm 0.01$ * | none | none |
| m1_none | -1485.9 | $-0.01 \pm 0.01$ * | none | none |
| m1_preg | -1498.2 | $-0.01 \pm 0.01$ * | none | none |
| m4_pweak | -1698.7 | $-0.01 \pm 0.01$ * | M.Gen.interval | PaperID |
| m4_none | -1701.9 | $-0.01 \pm 0.01$ * | M.Gen.interval | PaperID |
| m5_pfix | -1624.2 | $-0.01 \pm 0.01$ * | M.Gen.interval | PaperID + Kingdom + Binomen.rotl |
| m2_pweak | -1700.9 | $-0.01 \pm 0.01$ * | none | PaperID |
| m2_preg | -1694.9 | $-0.01 \pm 0.01$ * | none | PaperID |
| m2_none | -1697.1 | $-0.01 \pm 0.01$ * | none | PaperID |
| m6_pfix | -1624.5 | $-0.01 \pm 0.01$ * | M.Gen.interval | PaperID + Kingdom + Binomen.rotl<br>+ Marker.Type |
| m5_preg | -1641.9 | $-0.02 \pm 0.02$ * | M.Gen.interval | PaperID + Kingdom + Binomen.rotl |
| m5_pweak | -1709.2 | $-0.02 \pm 0.12$ ns | M.Gen.interval | PaperID + Kingdom + Binomen.rotl |
| m5_none | -1712.4 | $-0.03 \pm 0.11$ ns | M.Gen.interval | PaperID + Kingdom + Binomen.rotl |
| m6_preg | -1636.7 | $-0.03 \pm 0.03$ * | M.Gen.interval | PaperID + Kingdom + Binomen.rotl<br>+ Marker.Type |
| mshaw1_pfix | -1629.8 | $-0.04 \pm 0.01$ * | Z.Year.Midpoint + Z.Gen.interval | PaperID |
| mshaw1_preg | -1761.5 | $-0.04 \pm 0.02$ * | Z.Year.Midpoint + Z.Gen.interval | PaperID |
| mshaw1_none | -1762.0 | $-0.04 \pm 0.02$ * | Z.Year.Midpoint + Z.Gen.interval | PaperID |
| mshaw1_pweak | -1769.9 | $-0.04 \pm 0.02$ * | Z.Year.Midpoint + Z.Gen.interval | PaperID |

Table S 7: Log ratio diversity $\pi$ meta-regression model comparison. (continued)
| label | DIC | summary | fixed effects | random effects |
| --- | --- | --- | --- | --- |
| m6_pweak | -1778.1 | $-0.07 \pm 0.16$ ns | M.Gen.interval | PaperID + Kingdom + Binomen.rotl + Marker.Type |
| m6_none | -1775.9 | $-0.07 \pm 0.15$ ns | M.Gen.interval | PaperID + Kingdom + Binomen.rotl + Marker.Type |

#### *Allelic richness M* percent change

**Table S 8:**
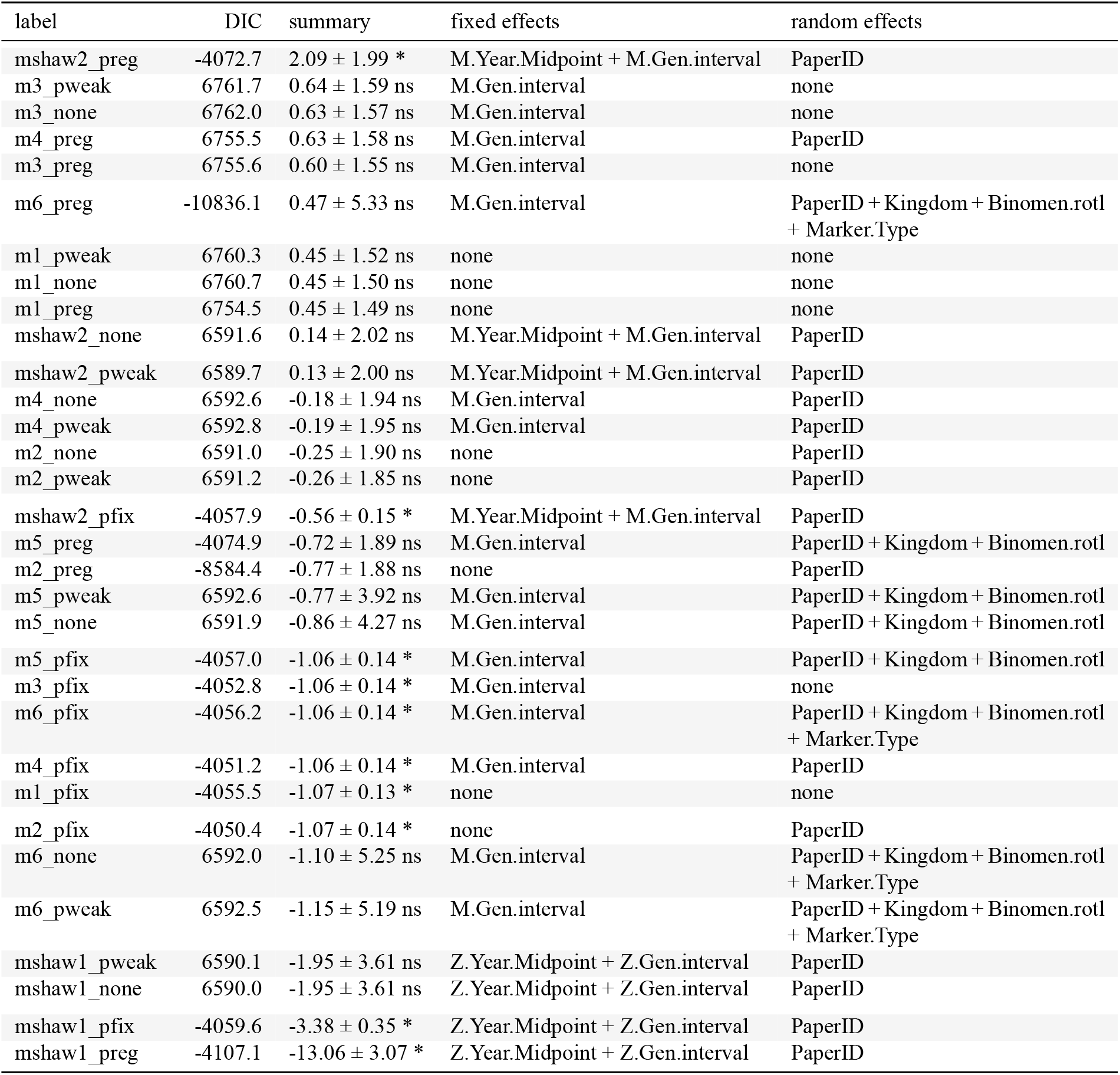
Percentage allelic richness M meta-regression model comparison.

**Allelic richness $M$ percent change**Table S 8: Percentage allelic richness $M$ meta-regression model comparison.
| label | DIC | summary | fixed effects | random effects |
| --- | --- | --- | --- | --- |
| mshaw2_preg | -4072.7 | $2.09 \pm 1.99$ * | M.Year.Midpoint + M.Gen.interval | PaperID |
| m3_pweak | 6761.7 | $0.64 \pm 1.59$ ns | M.Gen.interval | none |
| m3_none | 6762.0 | $0.63 \pm 1.57$ ns | M.Gen.interval | none |
| m4_preg | 6755.5 | $0.63 \pm 1.58$ ns | M.Gen.interval | PaperID |
| m3_preg | 6755.6 | $0.60 \pm 1.55$ ns | M.Gen.interval | none |
| m6_preg | -10836.1 | $0.47 \pm 5.33$ ns | M.Gen.interval | PaperID + Kingdom + Binomen.rotl + Marker.Type |
| m1_pweak | 6760.3 | $0.45 \pm 1.52$ ns | none | none |
| m1_none | 6760.7 | $0.45 \pm 1.50$ ns | none | none |
| m1_preg | 6754.5 | $0.45 \pm 1.49$ ns | none | none |
| mshaw2_none | 6591.6 | $0.14 \pm 2.02$ ns | M.Year.Midpoint + M.Gen.interval | PaperID |
| mshaw2_pweak | 6589.7 | $0.13 \pm 2.00$ ns | M.Year.Midpoint + M.Gen.interval | PaperID |
| m4_none | 6592.6 | $-0.18 \pm 1.94$ ns | M.Gen.interval | PaperID |
| m4_pweak | 6592.8 | $-0.19 \pm 1.95$ ns | M.Gen.interval | PaperID |
| m2_none | 6591.0 | $-0.25 \pm 1.90$ ns | none | PaperID |
| m2_pweak | 6591.2 | $-0.26 \pm 1.85$ ns | none | PaperID |
| mshaw2_pfix | -4057.9 | $-0.56 \pm 0.15$ * | M.Year.Midpoint + M.Gen.interval | PaperID |
| m5_preg | -4074.9 | $-0.72 \pm 1.89$ ns | M.Gen.interval | PaperID + Kingdom + Binomen.rotl |
| m2_preg | -8584.4 | $-0.77 \pm 1.88$ ns | none | PaperID |
| m5_pweak | 6592.6 | $-0.77 \pm 3.92$ ns | M.Gen.interval | PaperID + Kingdom + Binomen.rotl |
| m5_none | 6591.9 | $-0.86 \pm 4.27$ ns | M.Gen.interval | PaperID + Kingdom + Binomen.rotl |
| m5_pfix | -4057.0 | $-1.06 \pm 0.14$ * | M.Gen.interval | PaperID + Kingdom + Binomen.rotl |
| m3_pfix | -4052.8 | $-1.06 \pm 0.14$ * | M.Gen.interval | none |
| m6_pfix | -4056.2 | $-1.06 \pm 0.14$ * | M.Gen.interval | PaperID + Kingdom + Binomen.rotl + Marker.Type |
| m4_pfix | -4051.2 | $-1.06 \pm 0.14$ * | M.Gen.interval | PaperID |
| m1_pfix | -4055.5 | $-1.07 \pm 0.13$ * | none | none |
| m2_pfix | -4050.4 | $-1.07 \pm 0.14$ * | none | PaperID |
| m6_none | 6592.0 | $-1.10 \pm 5.25$ ns | M.Gen.interval | PaperID + Kingdom + Binomen.rotl + Marker.Type |
| m6_pweak | 6592.5 | $-1.15 \pm 5.19$ ns | M.Gen.interval | PaperID + Kingdom + Binomen.rotl + Marker.Type |
| mshaw1_pweak | 6590.1 | $-1.95 \pm 3.61$ ns | Z.Year.Midpoint + Z.Gen.interval | PaperID |
| mshaw1_none | 6590.0 | $-1.95 \pm 3.61$ ns | Z.Year.Midpoint + Z.Gen.interval | PaperID |
| mshaw1_pfix | -4059.6 | $-3.38 \pm 0.35$ * | Z.Year.Midpoint + Z.Gen.interval | PaperID |
| mshaw1_preg | -4107.1 | $-13.06 \pm 3.07$ * | Z.Year.Midpoint + Z.Gen.interval | PaperID |

#### *Allelic richness M* log ratio (geometric)

**Table S 9:** Log ratio diversity M meta-regression model comparison.

| label | DIC | summary | fixed effects | random effects |
| --- | --- | --- | --- | --- |
| mshaw2_pfix | -359.4 | $0.00 \pm 0.01$ ns | M.Year.Midpoint + M.Gen.interval | PaperID |
| m3_pfix | -355.9 | $0.00 \pm 0.01$ ns | M.Gen.interval | none |
| m4_pfix | -358.1 | $0.00 \pm 0.01$ ns | M.Gen.interval | PaperID |
| m3_preg | -310.4 | $-0.00 \pm 0.01$ ns | M.Gen.interval | none |
| m3_none | -305.7 | $-0.00 \pm 0.02$ ns | M.Gen.interval | none |
| m6_pfix | -358.3 | $-0.00 \pm 0.02$ ns | M.Gen.interval | PaperID + Kingdom + Binomen.rotl + Marker.Type |
| m3_pweak | -304.2 | $-0.00 \pm 0.02$ ns | M.Gen.interval | none |
| m5_pfix | -358.7 | $-0.00 \pm 0.01$ ns | M.Gen.interval | PaperID + Kingdom + Binomen.rotl |
| m1_pfix | -360.1 | $-0.00 \pm 0.01$ ns | none | none |
| m1_none | -306.3 | $-0.00 \pm 0.02$ ns | none | none |
| m2_pfix | -360.2 | $-0.00 \pm 0.01$ ns | none | PaperID |
| m1_preg | -312.7 | $-0.00 \pm 0.01$ ns | none | none |
| m1_pweak | -306.7 | $-0.00 \pm 0.02$ ns | none | none |
| mshaw2_preg | -561.5 | $-0.01 \pm 0.02$ ns | M.Year.Midpoint + M.Gen.interval | PaperID |
| mshaw2_none | -560.3 | $-0.01 \pm 0.02$ ns | M.Year.Midpoint + M.Gen.interval | PaperID |
| mshaw2_pweak | -561.3 | $-0.01 \pm 0.02$ ns | M.Year.Midpoint + M.Gen.interval | PaperID |
| m2_preg | -560.3 | $-0.01 \pm 0.02$ ns | none | PaperID |
| m4_preg | -563.2 | $-0.01 \pm 0.02$ ns | M.Gen.interval | PaperID |
| m2_pweak | -560.0 | $-0.01 \pm 0.02$ ns | none | PaperID |
| m4_pweak | -558.2 | $-0.01 \pm 0.02$ ns | M.Gen.interval | PaperID |
| m4_none | -555.9 | $-0.01 \pm 0.02$ ns | M.Gen.interval | PaperID |
| m2_none | -555.0 | $-0.01 \pm 0.02$ ns | none | PaperID |
| m5_preg | -559.4 | $-0.01 \pm 0.02$ ns | M.Gen.interval | PaperID + Kingdom + Binomen.rotl |
| m6_preg | -555.4 | $-0.01 \pm 0.03$ ns | M.Gen.interval | PaperID + Kingdom + Binomen.rotl + Marker.Type |
| m5_none | -556.6 | $-0.02 \pm 0.14$ ns | M.Gen.interval | PaperID + Kingdom + Binomen.rotl |
| mshaw1_pfix | -358.3 | $-0.02 \pm 0.03$ ns | Z.Year.Midpoint + Z.Gen.interval | PaperID |
| m5_pweak | -557.7 | $-0.02 \pm 0.15$ ns | M.Gen.interval | PaperID + Kingdom + Binomen.rotl |
| mshaw1_preg | -569.8 | $-0.03 \pm 0.03$ ns | Z.Year.Midpoint + Z.Gen.interval | PaperID |
| mshaw1_none | -560.0 | $-0.03 \pm 0.04$ ns | Z.Year.Midpoint + Z.Gen.interval | PaperID |
| mshaw1_pweak | -557.7 | $-0.03 \pm 0.04$ ns | Z.Year.Midpoint + Z.Gen.interval | PaperID |
| m6_none | -560.0 | $-0.04 \pm 0.15$ ns | M.Gen.interval | PaperID + Kingdom + Binomen.rotl + Marker.Type |
| m6_pweak | -561.0 | $-0.04 \pm 0.16$ ns | M.Gen.interval | PaperID + Kingdom + Binomen.rotl + Marker.Type |

#### Test per-study statistical differences: are there more or less positive trends?

Is the distribution of Hedge’s g significantly different from zero for a given study? To test this, we follow a t-statistic distribution, which is the expected for Hedge’s g. In addition we also conduct the test with a Gaussian distribution (which may be more akin to the meta-regression modeling, but this Normal distribution causes less significant results than t-distribution because is wider and thus more conservative). Figure S4 shows all studies ranked by −*log*_10_(*p*) of Hedge’s g test per study, which has a substantial inflation. Figure 3 shows the inflation of P-values assuming Hedges’g follow a t-distribution, making our detection of significant effects anti-conservative.

Distribution of positive and negative π div trajectory between two time points. Using Hedge’s g for significance using two sided test and family-wise Bonferroni *P*-value correction 0.05/number of tests. The number of significant tests is as follows: Non-significant 823. Declining 50. Increasing 43.

- *For diversity π* Number significant trends (i.e. either positive or negative) = 93 of total 916 points. Percentage of significant trends (i.e. either positive or negative) = 10%
- *For richness M* Number significant trends (i.e. either positive or negative) = 112 of total 785 points. Percentage of significant trends (i.e. either positive or negative) = 14%

Are the number of significantly negative (decline of genetic diversity) and positive (increase in genetic diversity) more or less common? We conduct a Binomial test of enrichment of significant tests that are negative vs positive:

- *For diversity π* Prob. success (i.e. positive vs negative) = 0.4623656 P value = 0.5340599 There are also 46 *P*-values=0 (5.02% of dataset). The cases of *P*-value=0 are roughly equal in both directions, 24 decreasing, 22 increasing.
- *For richness M* Prob. success (i.e. positive vs negative) = 0.4196429 P value = 0.1077823 There are also 31 *P*-values=0 (3.95% of dataset). The cases of *P*-value=0 are roughly equal in both directions, 17 decreasing, 14 increasing.

**Figure S 3:**
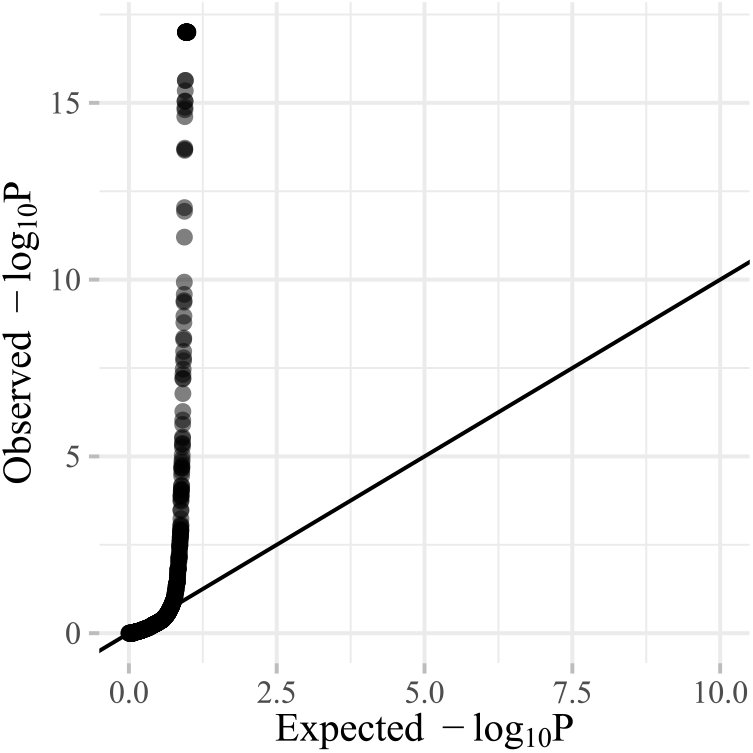
P-value distribution in QQ plot.

We also test this with all the data points from Shaw et al (4023) to make sure this is not a subset effect:

- *For all dataset* Prob. success (i.e. positive vs negative) = 0.5099237 P value = 0.6391896

In addition, I compute the same binomial test without test statistics. Simply, count how many studies have Hedges’ g (effect size) that is negative (2095) out of all the dataset (4023) and ask whether this is different from 50/50. There is no significant divergence from 50/50, with estimate 0.5207557 and P-value=0.995964.

**Figure S 4:**
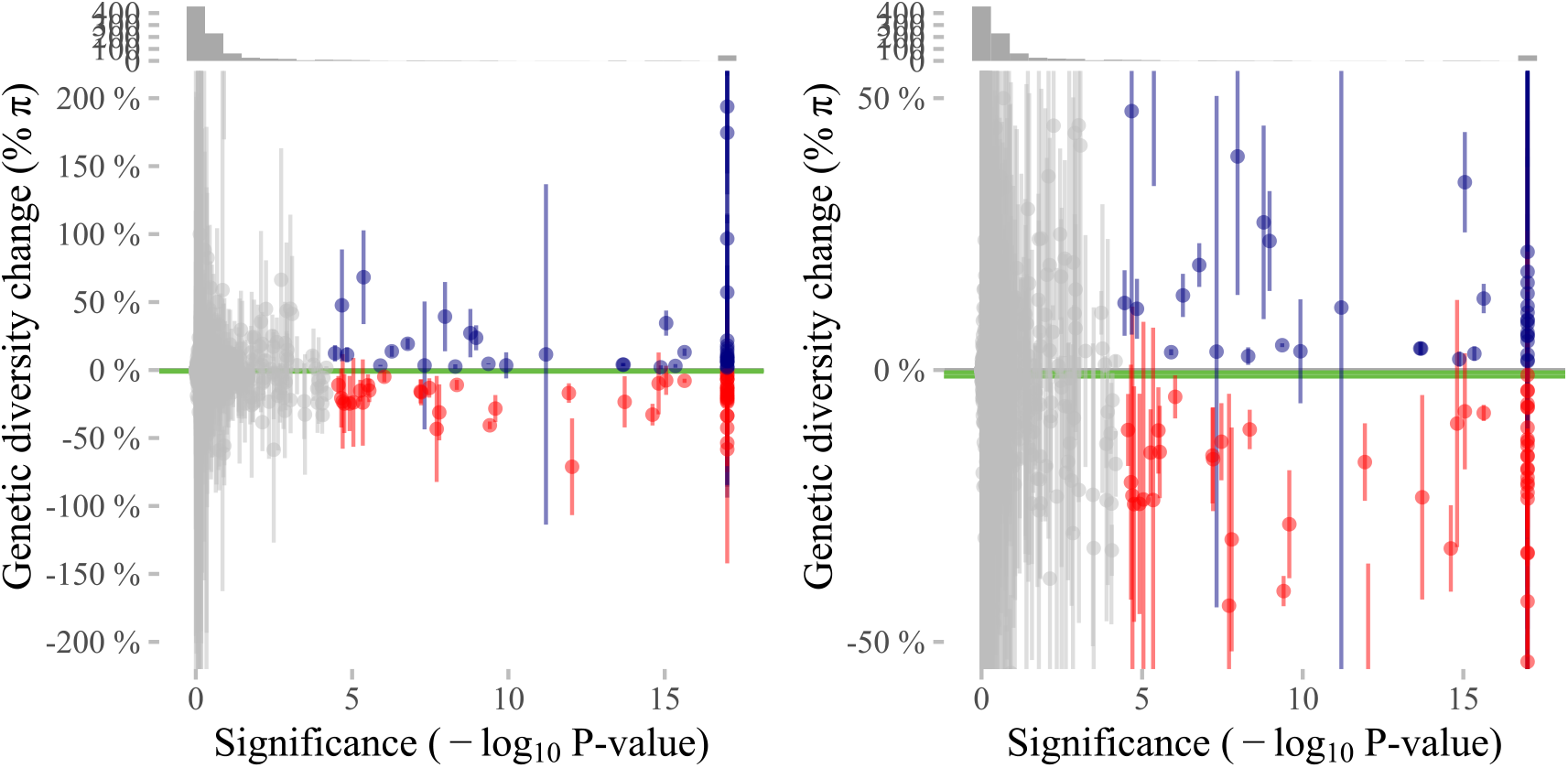
Relationship between raw values of diversity change π and Hedge’s g significance.

**Figure S 5:**
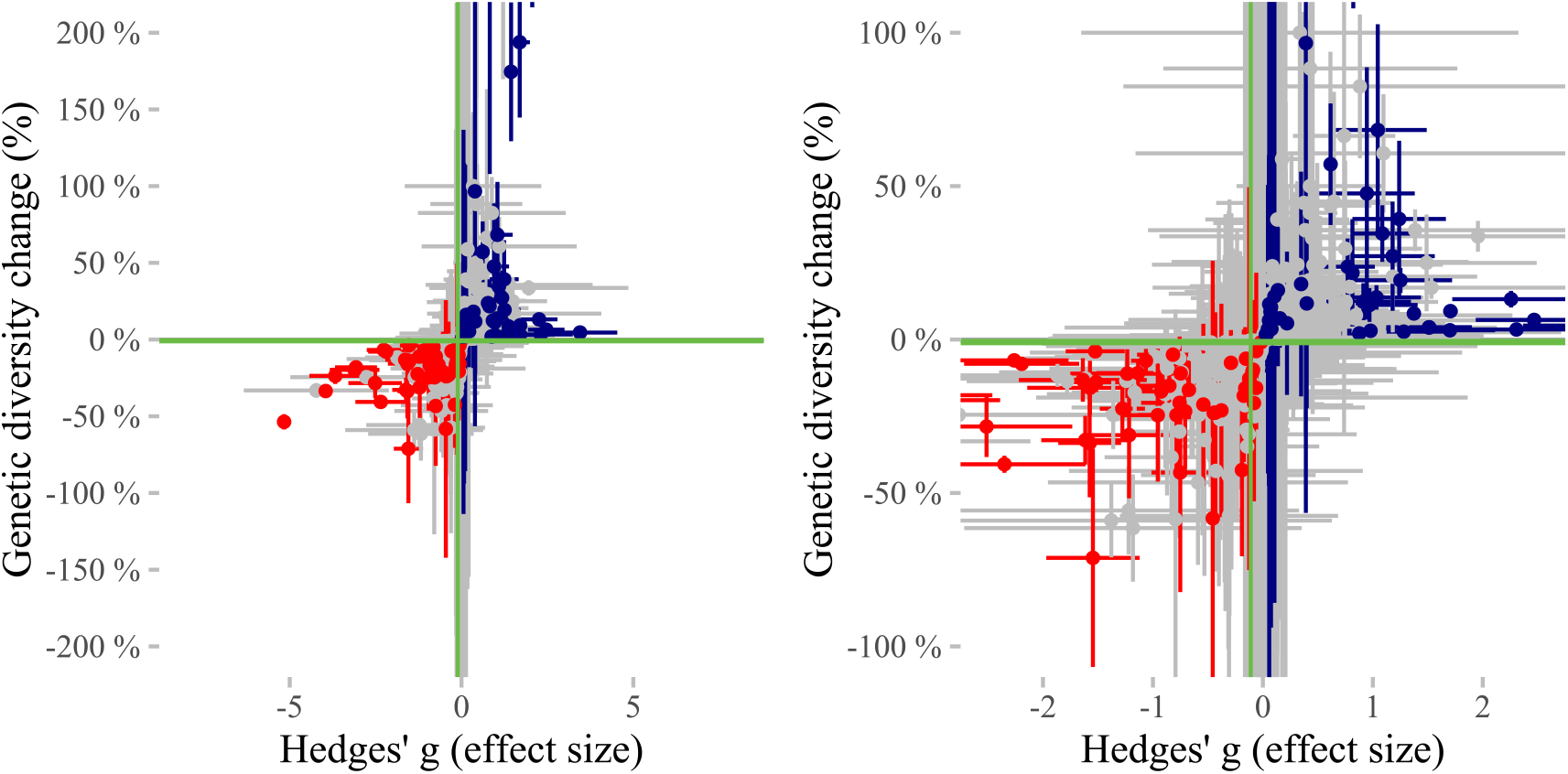
Relationship between raw values of diversity change π and Hedge’s g values.

#### Test relationship with Red List

**Figure S 6:**
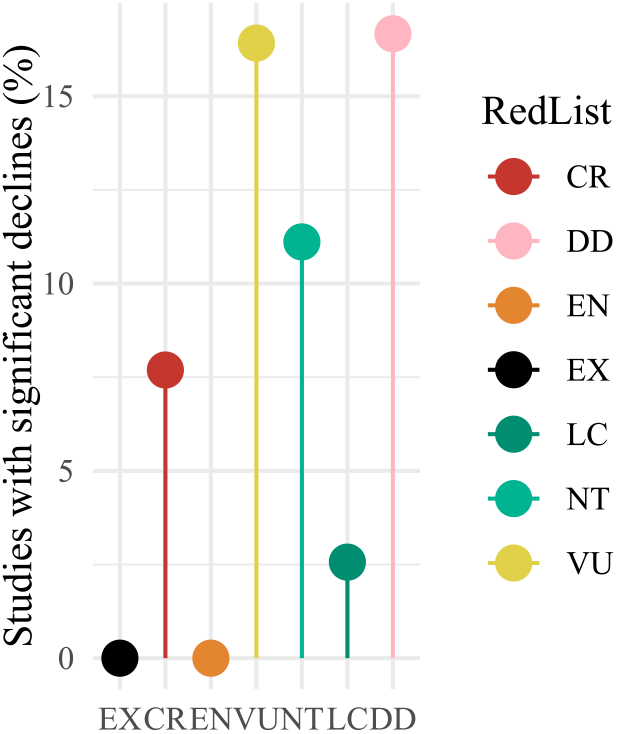
Significant decline trends and Red List categories.

The number of significant decline cases in π (50), was significantly enriched with species in the Red List (all but least concern [LC] or data deficient [DD], 475, Fisher’s test *P* =0.0012471, Odds=2.7783496) (Fig. 6). The number of significant genetic diversity increase cases too (Fisher’s test *P* =0.1606518, Odds=1.6000509). This was mostly driven by vulnerable species (6).

#### Test differences with increasing time

The number of significant decline cases in π (50), were marginally significantly enriched with high number of generation interval (>30 generations # studies= 37, Fisher’s test *P* =0.0457039, Odds=2.8906597) (Fig. 7). This was non-significant if excluding positive trends (which may not be realistic) (*P* =0.0543964, Odds 2.7419411). Only focusing on studies with increasing genetic diversity trends showed also no enrichment (Fisher’s test *P* =0.411707, Odds=0).

#### Test differences with increasing data size

There was a significant enrichment of trajectories of π% decline in studies with number of loci above 10 (Odd=15.8647827, Fisher’s test *P* =1.2975938 × 10^−12^) as well as π% increases (Odd=7.5969563, Fisher’s test *P* =1.51457 × 10^−8^).

There was not a significant enrichment of trajectories of π% decline in studies with >100 individuals sampled (Odd=0.1889661, Fisher’s test *P* =0.0774391) nor π% increases (Odd=0.9965217, Fisher’s test *P* =1).

#### Test differences with decreasing noise

It has become clear that the very extreme values of genetic diversity increases and decreases are owed to high standard error. A good rule of thumb is that standard error < 5% are accurate measurements, while >5% are highly uncertain, as the measurements we are aiming to achieve are in the same or smaller order of magnitude. We then look at enrichment of significant values in the more certain studies. I find a marginal enrichment of significant decline diversity cases π and low error measurements (Odd=1.8501645, Fisher’s test *P* =0.0443237) but not with significantly increasing π studies (Odd=1.7378191, Fisher’s test *P* =0.0966327).

#### Test differences with domesticated vs not species

Finally, hat the domesticated/pathogens/pest speices not being removed, we would find an upward bias in genetic diversity change as there is an enrichment of domesticated species and significant positive trends in the original data (Odds= 1.8437971, Odds= 3.9582345 × 10^−6^).

**Figure S 7:**
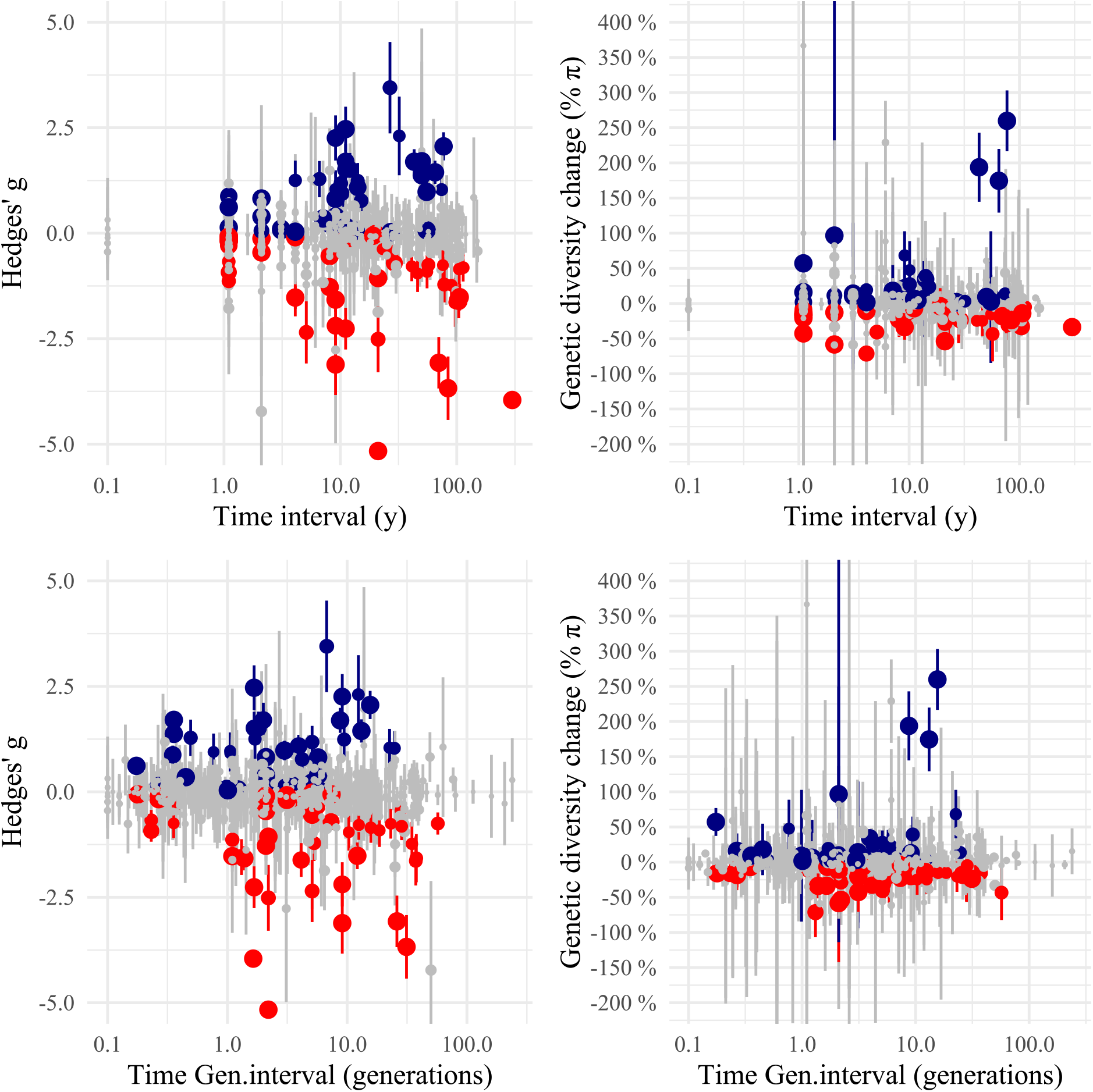
Relationship between genetic changes and length of time interval.

**Figure S 8:**
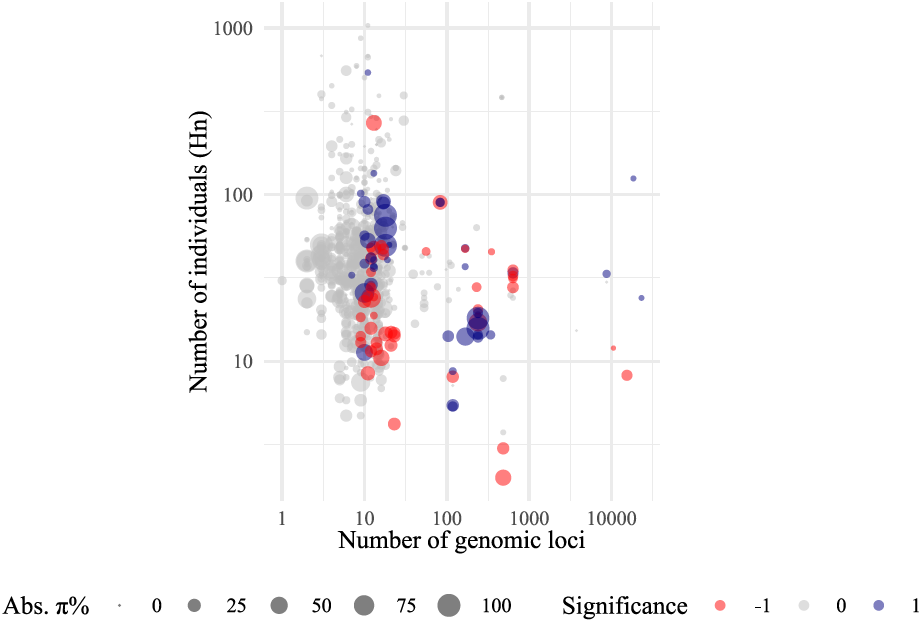
Sample size (loci and individuals) and raw changes in π% genetic diversity.

**Figure S 9:**
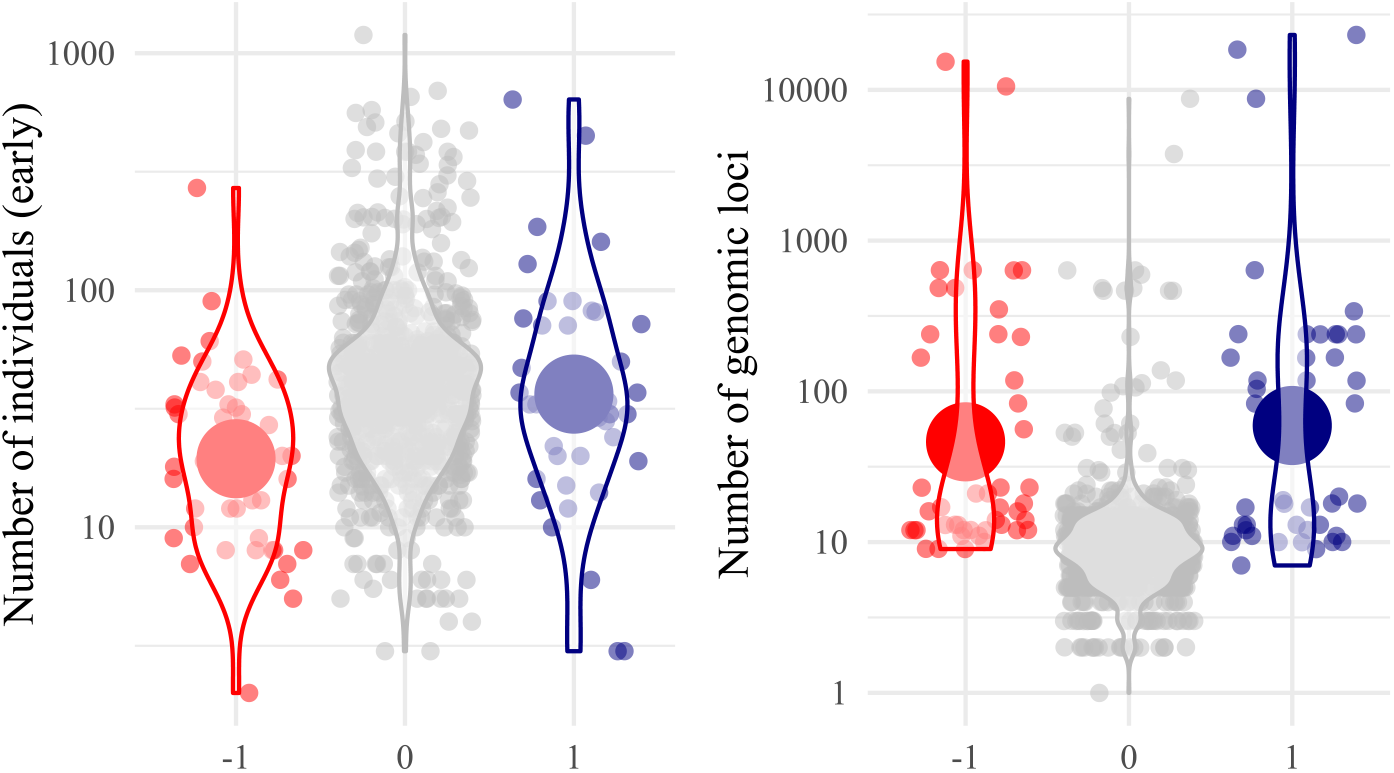
Sample numbers across past and present studies.

**Figure S 10:**
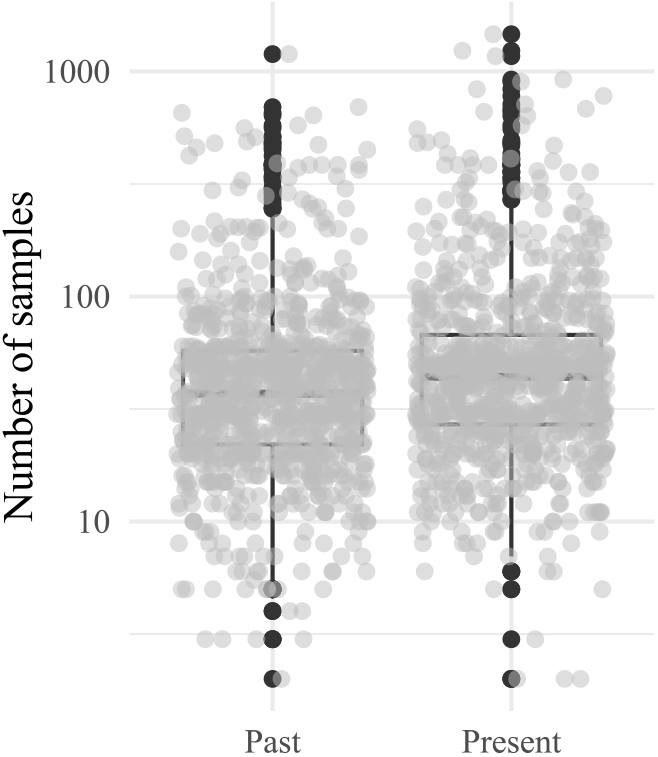
Sample and marker numbers across studies

**Figure S 11:**
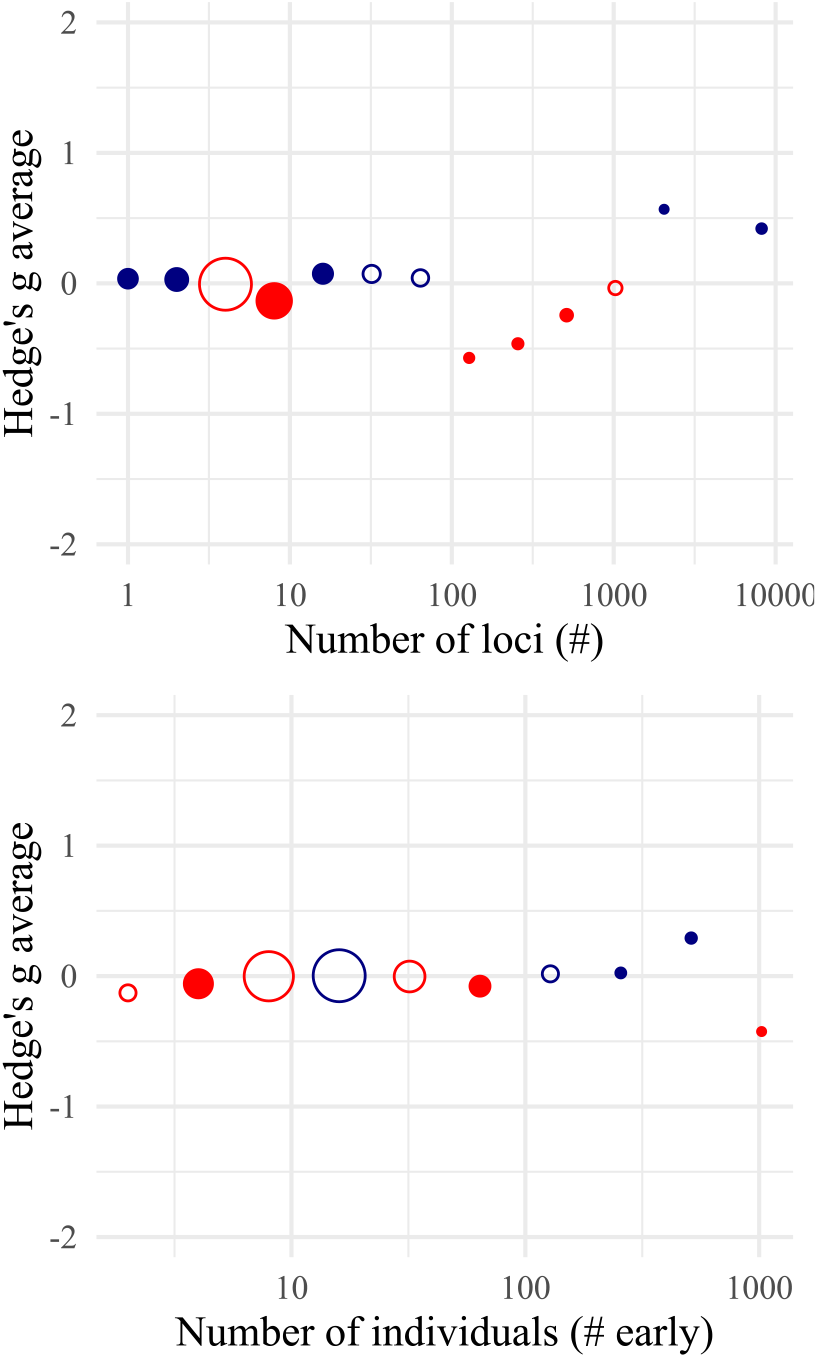
Average diversity change of different data sizes.

**Figure S 12:**
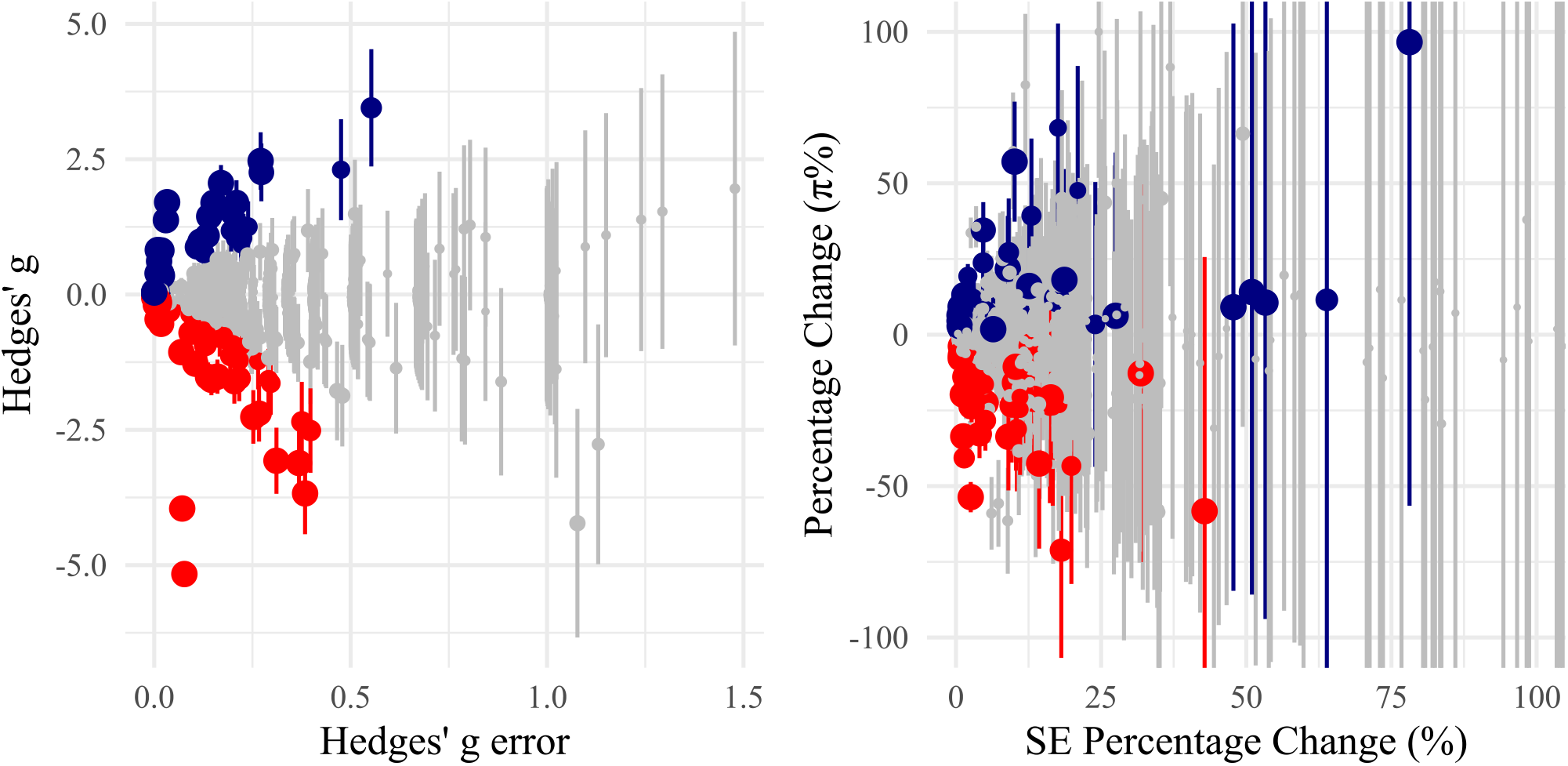
Relationship between raw values, significance and noise

### Text S3: Test within-species consistency

There were repeated measures per species in the dataset. For instance, for π there were 916 studies, but withihn these there were a total of 323 unique species.

We hypothesize that if the observed decline are robus they should occur in the same direction within the same species We then conduct a Binomial test of species with multiple significant observations, asking whether they tend to have more positive or more negative significant values.

**Figure S 13:**
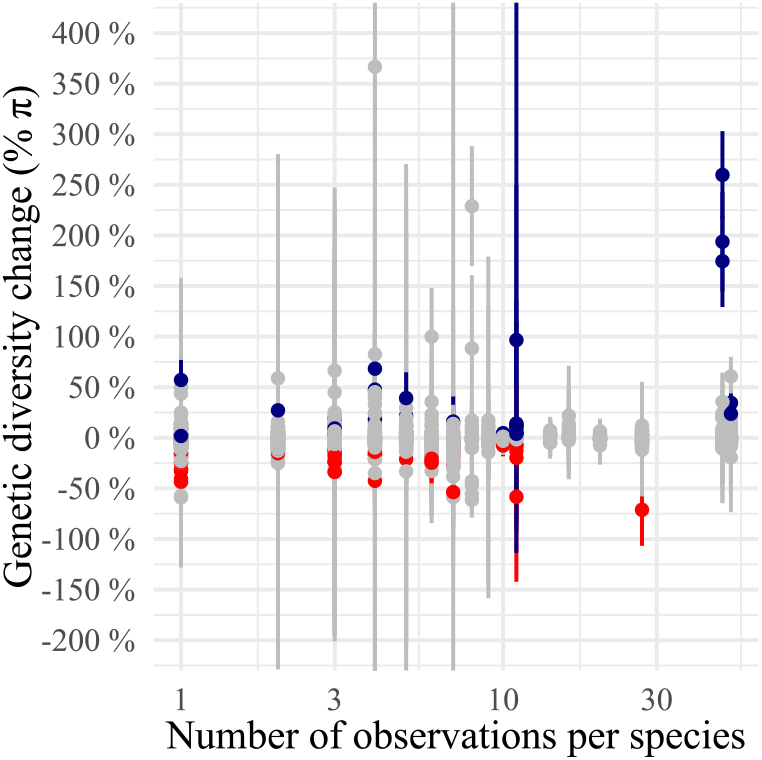
Cases of species with multiple observations.

**Figure S 14:**
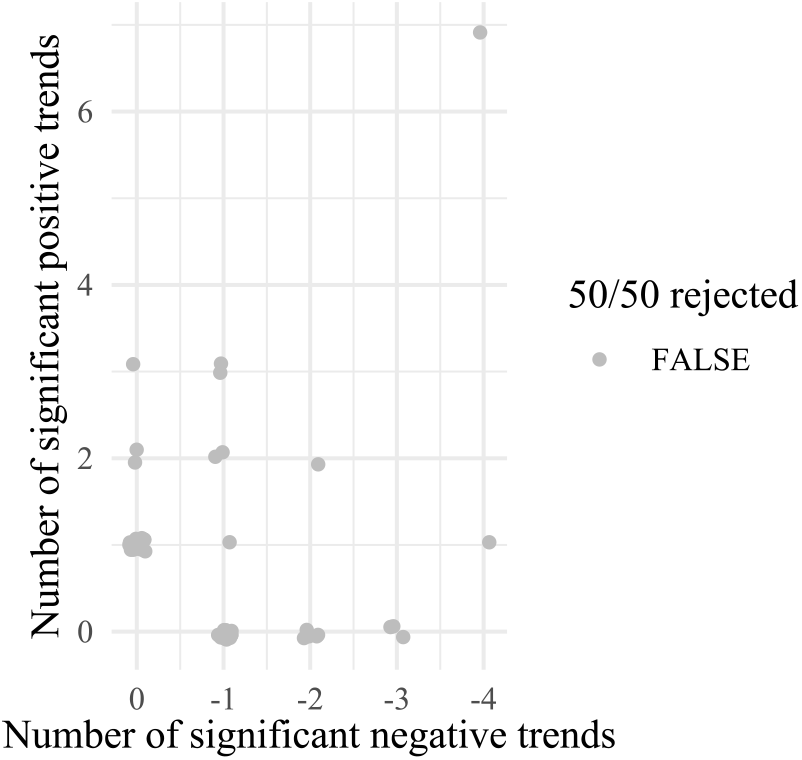
Positive and negative cases of species with multiple observations.

Of 50 species with multiple populations sampled and significant trends, none had a ratio of negative trends (decline) vs positive trend (increase) that was significant under a Binomial test of expected 50/50% probability.

### Text S4: Interpretation of changes under an evolutionary population genetic model of species contraction

Below we explore what % genetic diversity changes (non-significant or significant) could mean in terms of population contraction models of different scenarios (we assume positive changes are zero, as they are treated separately, see Text S5). Equations are derived in the **Mathematical Appendix**.

Some key notation:

- π nucleotide diversity or expected heterozygosity (which are identical under a biallelic loci condition).
- *M* allelic richness, or segregating sites, or number of variable positions, or number of mutations in a strand of DNA. Note we use *M* instead of more standard *S* or *A* to avoid confusions with species richness from ecology, and also to avoid confusions with area of habitat.
- *N*_0_ the population size at some past time. We may also use *N* as this is the long-term equilibrium size.
- *N*_*t*_ the population size at a given time *t*. Because we mostly deal with two time points (0 and 1), we often use *N*_1_ the population size instantaneous reduction from human impact or contraction.
- *A*_0_ the habitat area of a species with multiple populations (useful to use area to avoid defining discrete entities).
- *A*_1_ the habitat area of a species reduced from human impact or contraction.
- *X*_π_ the fraction loss of genetic diversity π. Intuitive % of genetic diversity loss when *X*_π_ × 100.
- Same for *X*_*M*_ the fraction loss of allelic richness *M*.

### BIOLOGICAL SAMPLING – *null scenario*

***Average genetic diversity*** π The high noise is likely due to the small sampling sizes:

Present sampling median n0=48

Present sampling median n1=52

Number of markers median L=9

Taking biological sampling into account, we found 22 studies out of 306 with changes in measured genetic diversity (−18%) beyond the 5% tail of the null distribution from subsampling (−8.4%). This corresponds to an excess of −10%.

Bootstrapping 1000 times one simulation per paper gives a range of average π% change= −1.1 - 6%.

The reason observations are very noisy is because the median sampling effort in Shaw is *n*_0_=37, *n*_1_=44, and the median number of loci is *L*=10. Fig. 18 shows the results of standard deviation of percent diversity change (%π) in a population of constant size sampled two times, in a grid of values from 5 individuals to 10,000 individuals, and a number of markers 1 to 1M. The median values of Shaw are marked in green, and indicate that the median error rate is over 10%, just as we see in the data.

Based on Fig. S17, the expected error for 50% of the Shaw dataset is a minimum of 32.6933738%, and for 75% percentile of the dataset a minimum of 11.3687606%.

### *Allelic richness M* TBD

**Figure S 15:**
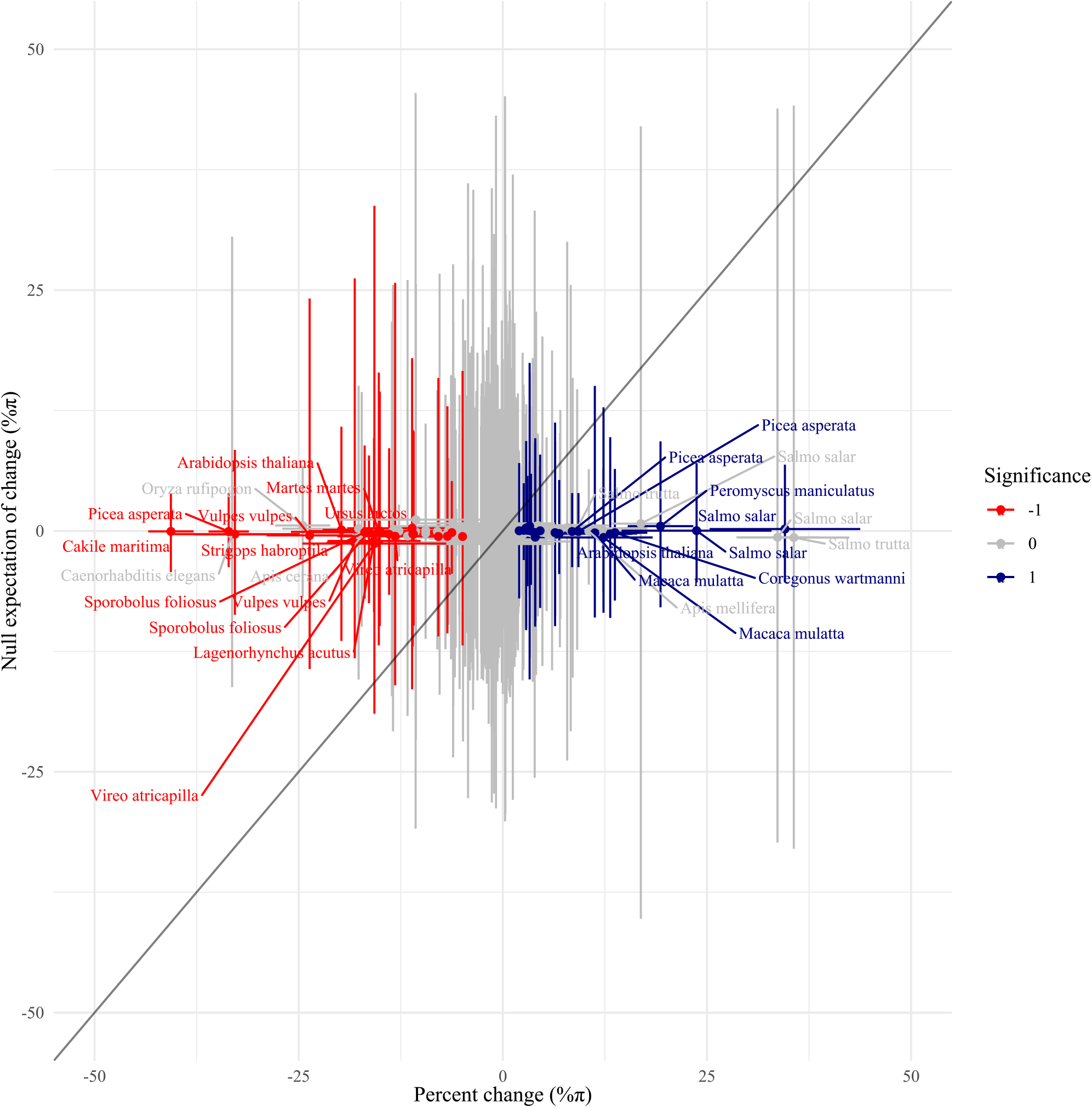
Ranges of expected changes of diversity π% with biological sampling

**Figure S 16:**
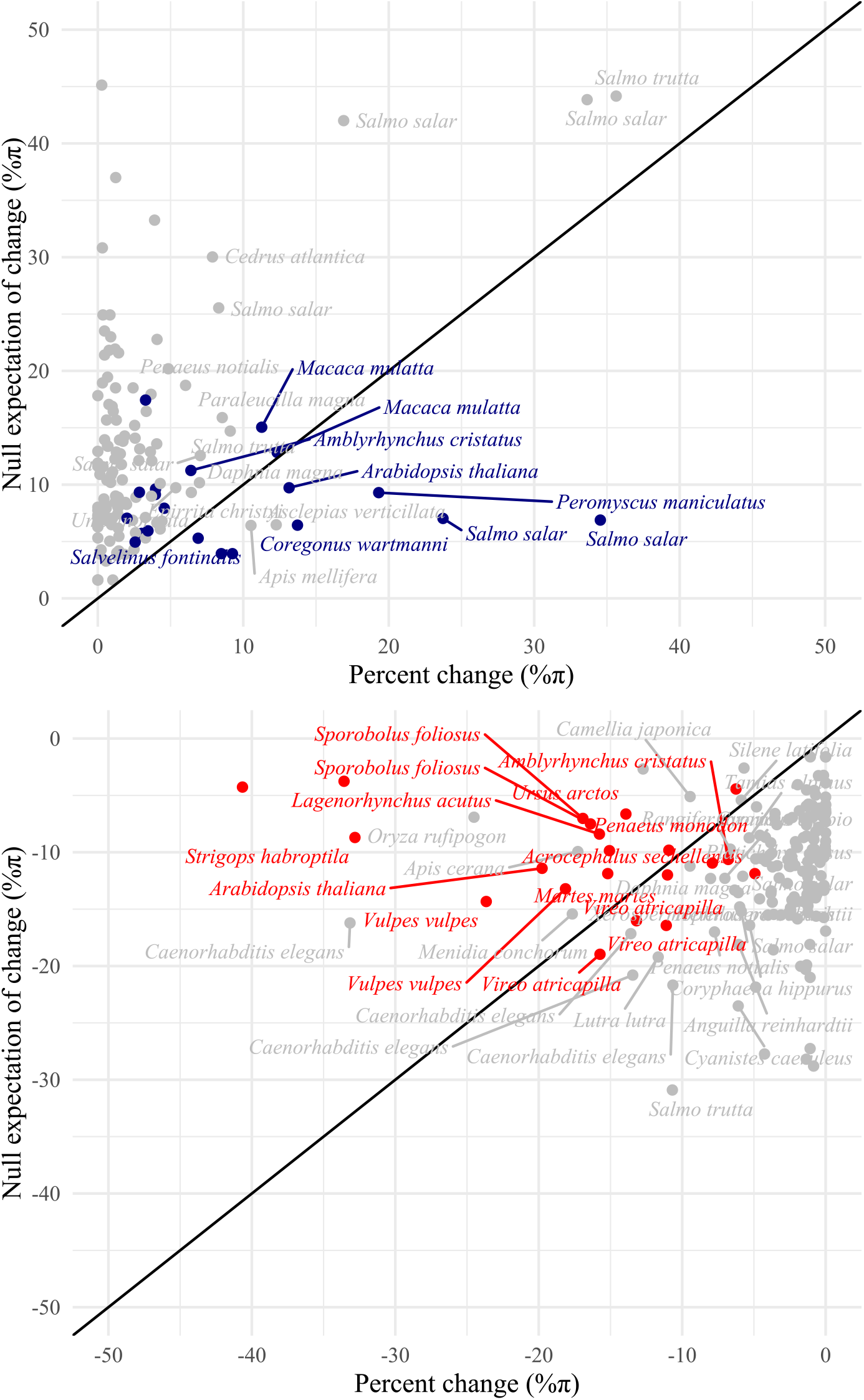
Expected maximum changes of diversity π% with biological sampling

**Figure S 17:**
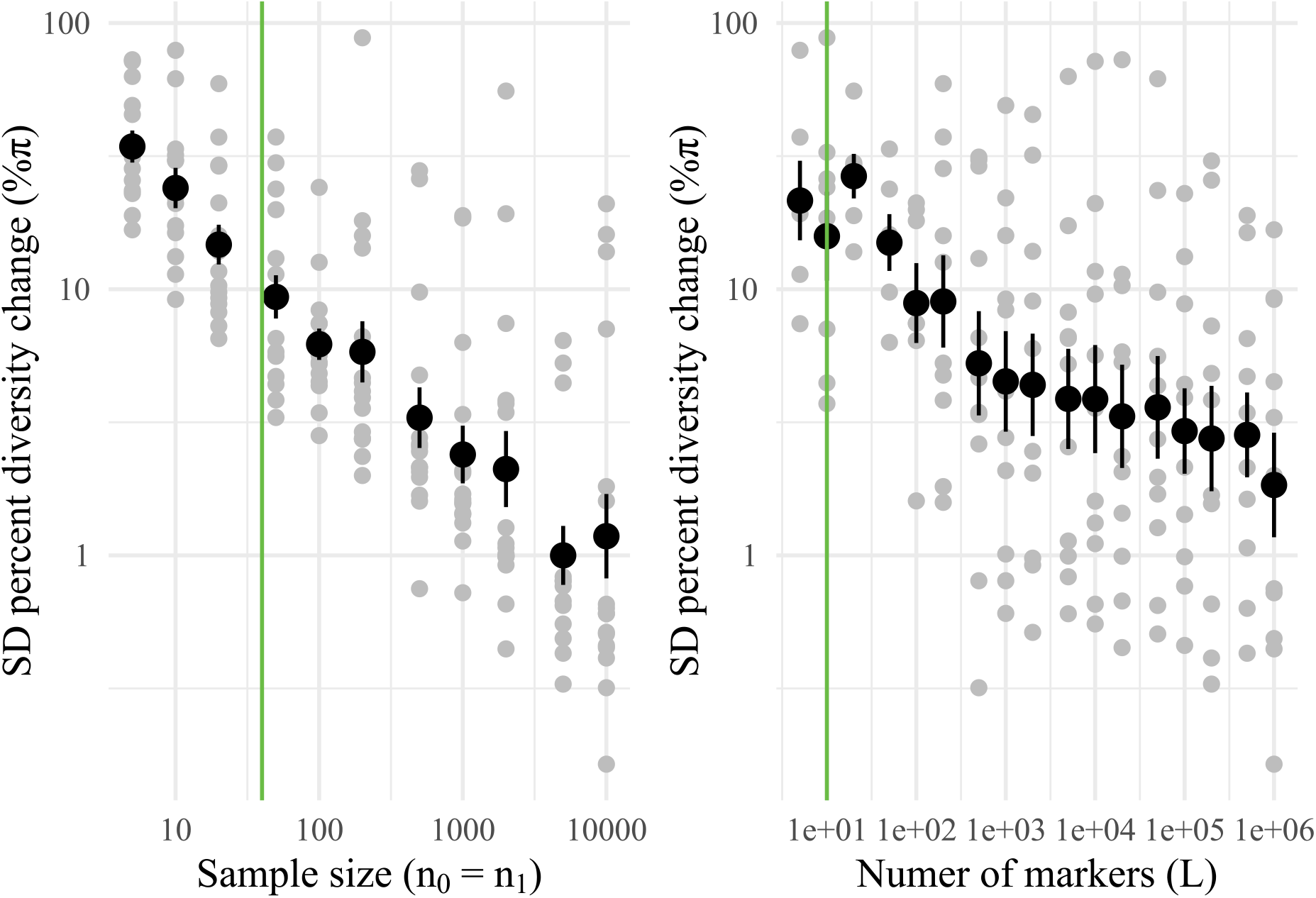
Expected noise in percent diversity change π% with different ssample sizes

### SUBSAMPLE Single population, immediate – *scenario 1a*

#### *Average genetic diversity* π

Based on population evolutionary genetic theory, in the **Mathematical Appendix**, we show that the expected relative π% change (or the fraction change, *X*_π_, which is more mathematically convenient) from a reduction of *N*_0_ to *N*_1_ within a generation is null. The only reason genetic diversity may change as a subsample of a population is due to sampling noise or if for some reason the remaining fraction of the population is more related to each other. This affects a fraction of the dataset: 295 observations, 32%. The results in Fig. 18, corroborate the classic observation that substantial losses (∼5%) of genetic diversity are not expected unless the bottleneck in a species is «100 individuals. NB. Fig. 18: A jitter of 0.1N units was added for visualization. Min value of bottleneck truncated to 10M. Positive trend % of genetic diversity collapsed to 0%.

**Figure S 18:**
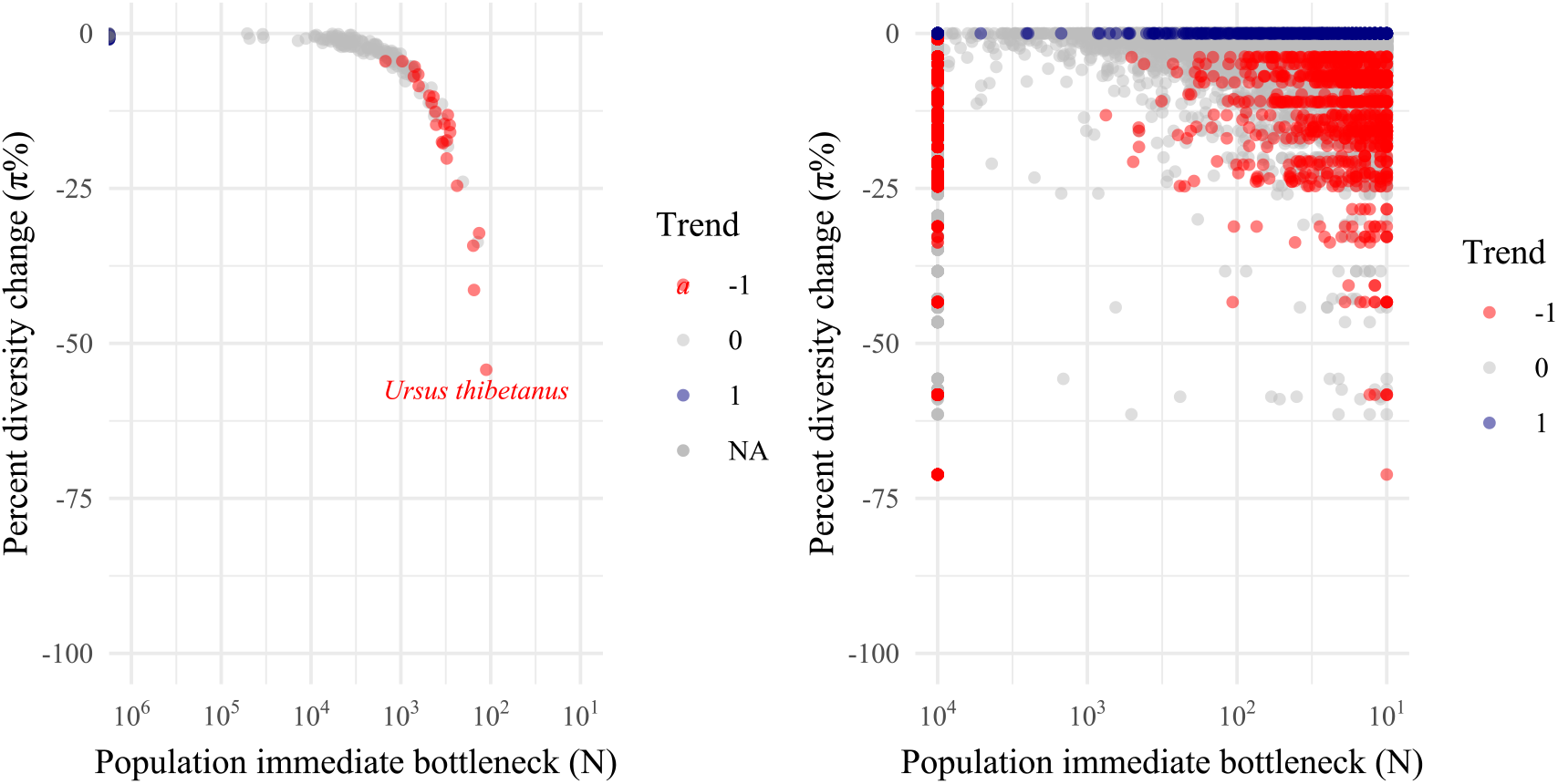
Expected immediate population bottleneck given observed π% with noise

*No noise* Median = 330 individuals.

Mean = 436 individuals.

IQR = 270 - 501 individuals.

Range = 93 - 1307 individuals.

*Noise* Median = 4 individuals.

Mean = 10 individuals.

IQR = 2 - 9 individuals.

Range = 1 - 752 individuals.

#### Allelic richnes M

The loss of alleles *M* follows under the coalescent a function of: 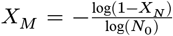. In the case of alleles *M* the loss not only depends on the fraction loss of individuals *X*_*N*_, but also the starting population sizes *N*_0_ (this is unknown, so Fig. **??** shows trajectories with several *N*_0_). NB Fig. **??** A jitter of 1N units was added for visualization. Positive trend % of genetic diversity collapsed to 0%. The patter observed of four trajectories correspond to inferences assuming *N*_0_= 100, 1000, 10000, 1000000 starting individuals prior the bottleneck (respectively left to right).

Using the results assuming a starting size of *N*_0_=10000 individuals. For allelic richness *M* the loss is faster than for π diversity, as expected.

**Figure S 19:**
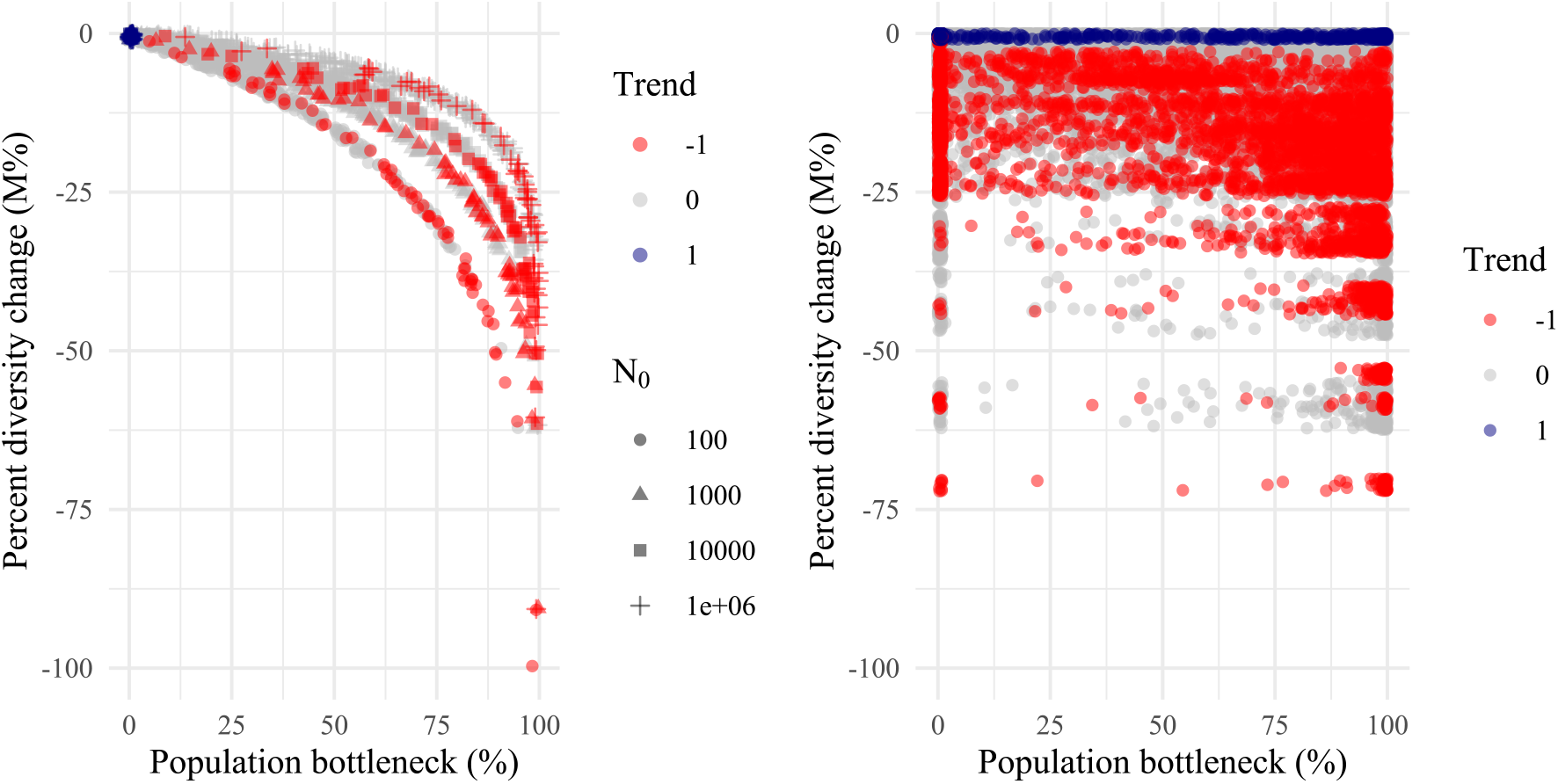
Expected immediate population bottleneck given observed M%.

*No noise* (assume the case of *N*_0_=10000) Median = NA% of original population

Mean = NaN% of original population

IQR = NA - NA% of original population

IQR = Inf - −Inf% of original population

*Noise* (assume the case of *N*_0_=10000) Median = 80% of original population

Mean = 62% of original population

IQR = 23 - 96% of original population

IQR = 0 - 100% of original population

### TIME Single population, drift over time – *scenario 1b*

**Figure S 20:**
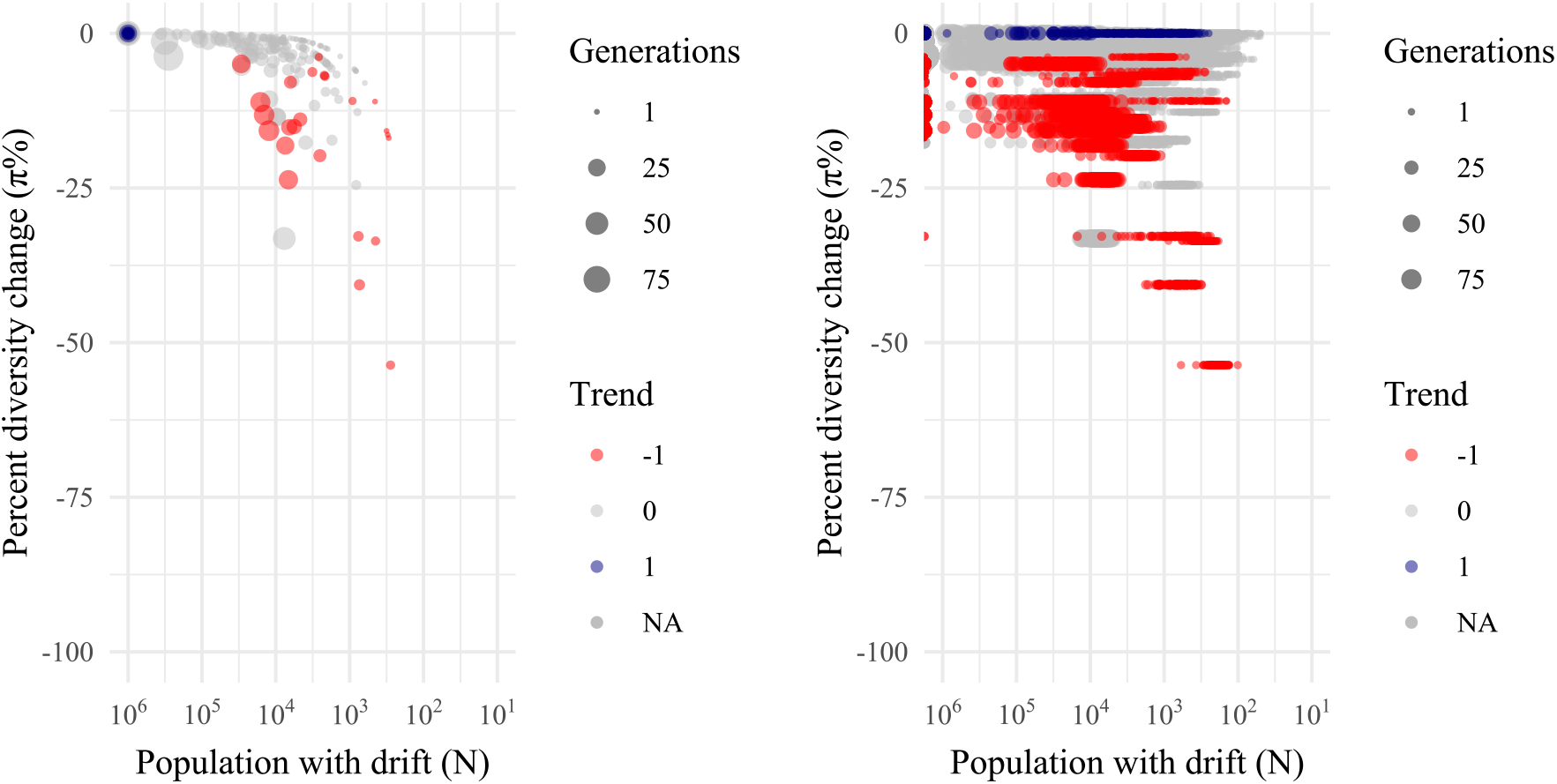
Expected population bottleneck and drift given observed π%.

#### *Average genetic diversity* π

NB. Positive trend % of genetic diversity collapsed to 0%.

Number of generations truncated to 100+ for visualization purposes.

Number of population size needed to explain loss truncated to 1M.

*No noise*

Median = 2570 individuals.

Mean = 5292 individuals.

IQR = 666 - 6623 individuals.

Ranges = 279 - 29401 individuals.

*Noise*

Median = 2624individuals.

Mean = Infindividuals.

IQR = 426 - 9195individuals.

Ranges = 0.5 - Infindividuals.

#### Allelic richnes M

Population bottleneck truncated up to 10^5^ for visualization Positive trend % of genetic diversity collapsed to 0%.

Starting population also needs to be assumed, *N*_0_ = 10^6^, which is a conservative value (as a smaller starting opulation is assumed, the bottleenck predicted is even larger)

Expected instantaneous population bottleneck percentage for studies with significan decline of *M* :

**Figure S 21:**
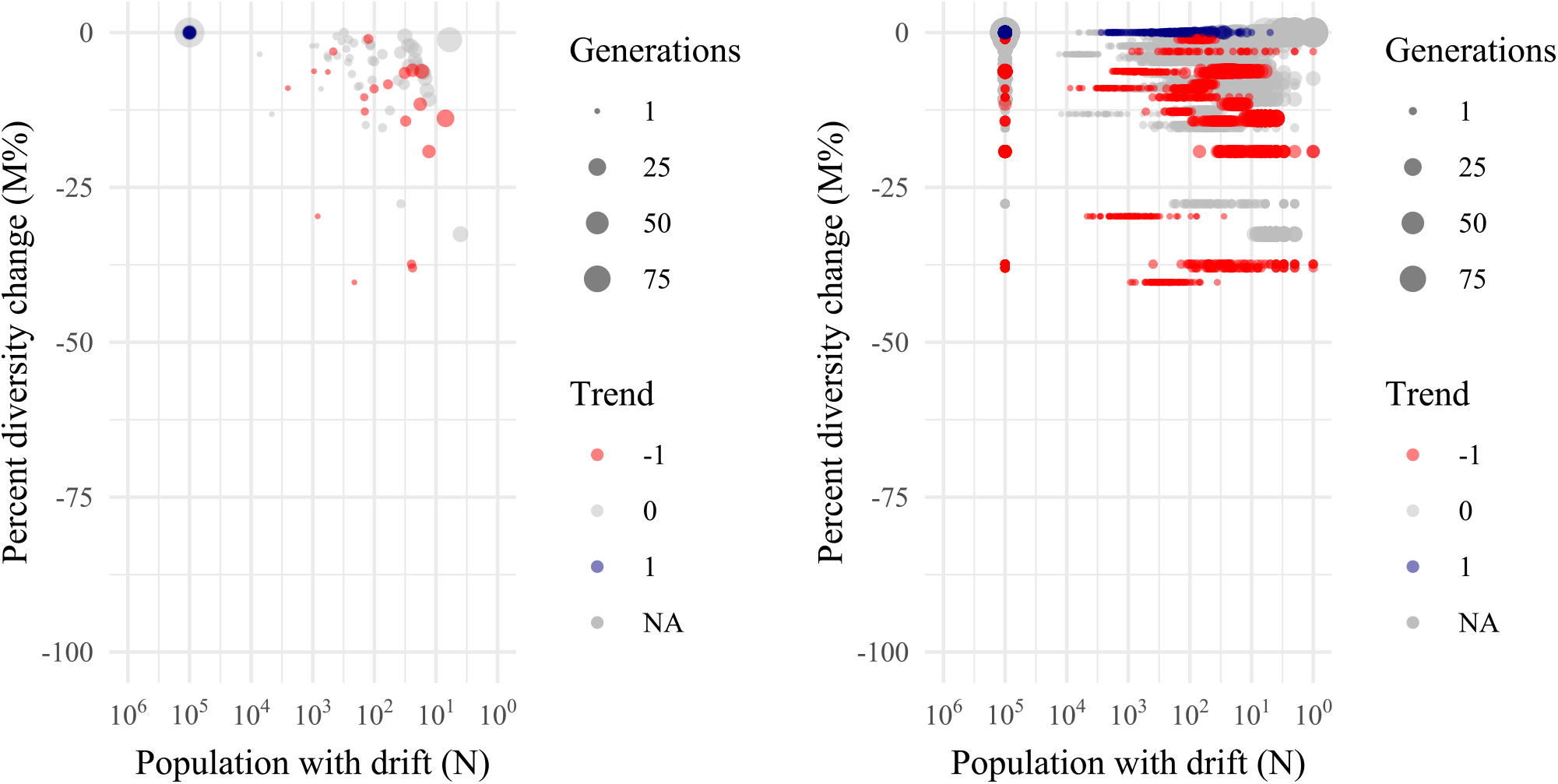
Expected immediate population bottleneck given observed M%.

Median = 72% of original population

IQR = 22 - 486

### SPACE. Spatial meta-populations, immediate – *scenario 2a*

**Figure S 22:**
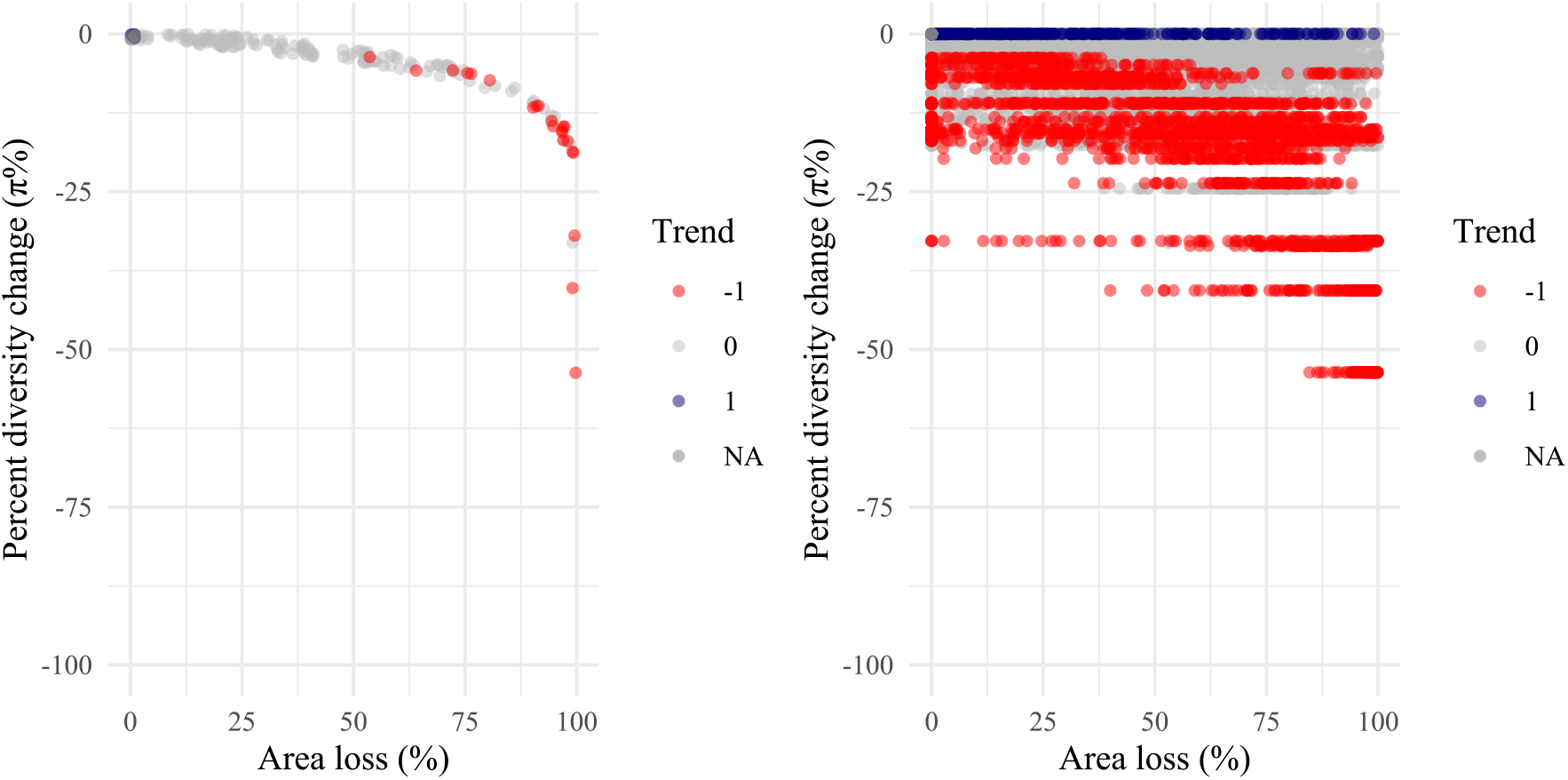
Expected immediate geographic area loss given observed π%.

#### *Average genetic diversity* π

NB. A jitter of 1% area units was added for visualization. Positive trend % of genetic diversity collapsed to 0%.

*No noise*

Inverse modeling of instantaneous *M* richness loss with spatial contraction:

Median = 96% area

Mean = 90% area

IQR = 88 - 98% area

Range = 54 - 100% area

*Noise*

Inverse modeling of instantaneous *M* richness loss with spatial contraction:

Median = 55% area

Mean = 51% area

IQR = 25 - 79% area

Range = 0 - 100% area

Expected instantaneous area contraction percentage for studies with significan decline of π :

Median = 55% of original range

IQR = 25 - 79

#### Allelic richnes M

Positive trend % of genetic diversity collapsed to 0%.

The above reconstructions take at face value the average genetic diversity change, but as seen in Text S2 there is much noise in these estimates. A more true visualization would re-generate each observation 100 times following the observed error in the quantity.

Expected instantaneous area contraction percentage for studies with significan decline of *M* :

**Figure S 23:**
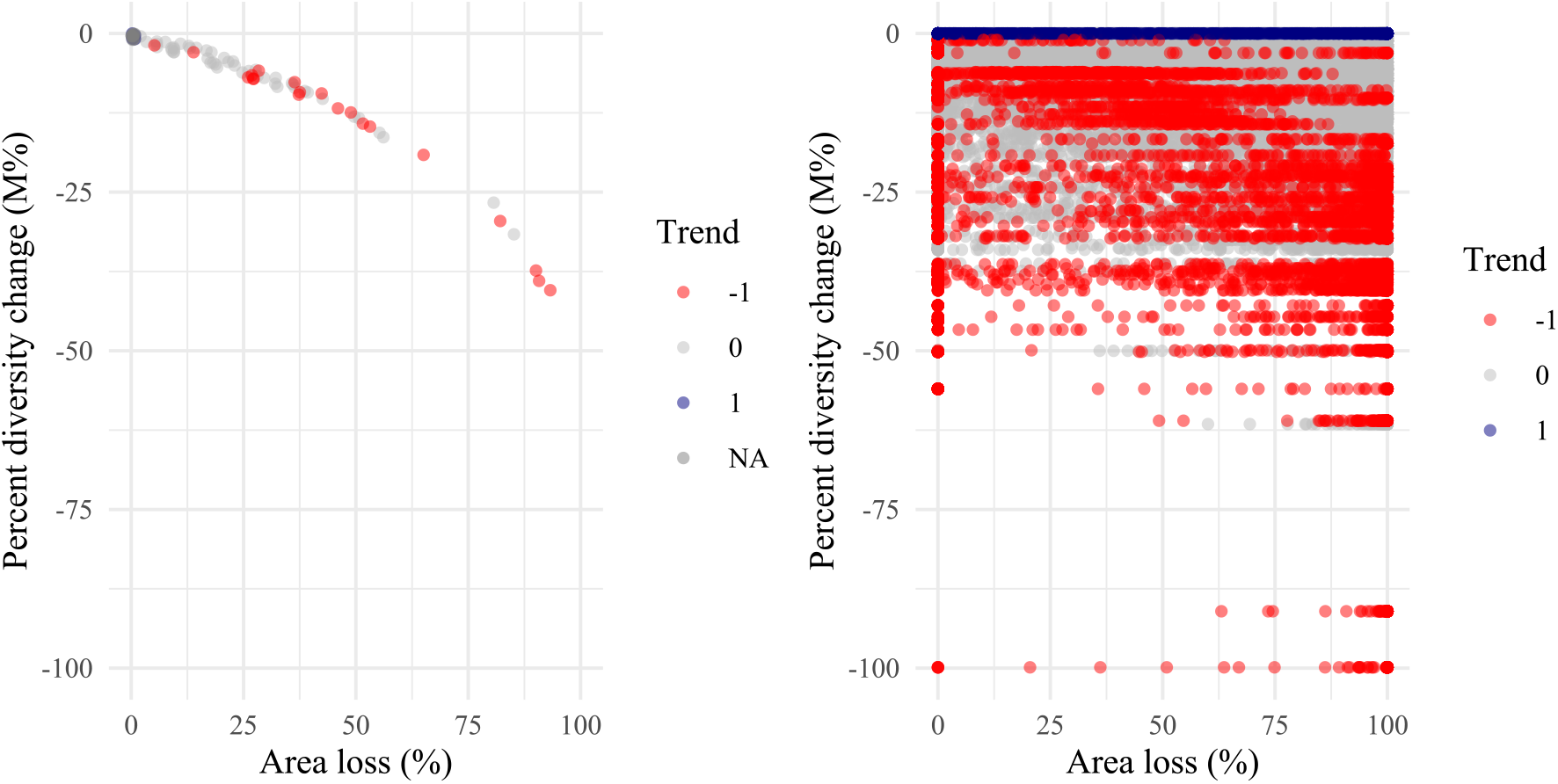
Expected immediate geographic area loss given observed M%.

Median = 61% of original range.

IQR = 0 - 97.

### SPACE & TIME. Spatial meta-populations via WFmoments, short drift – *scenario 2b*

Both spatial contraction and short drift. This is the most complicated scenario as we do not have a direct simple mathematical method. Instead, we have pre-computed scenarios of a solved numerical method we call WFmoments, that can predict the non-equilibrium genetic diversity of any meta-population system over time. We also only have solutions for average genetic diversity π. In this approach, then we created a number of loss area scenarios in a system of 100 demes in a 2D lattice, losing demes in an edge contraction manner. We then inverse model the area loss corresponding to the matched π change simulated and the observed π changes. We can use multiple functions, although they give qualitatively similar results. We decide for a Gaussian kernel, where the distance between the observed and simulated π changes exponentially decreases.

By definition, meta-population dynamics will be most relevant to species with multiple populations, and thus may not be as affected by single population dynamics (i.e. with a number of populations in the landscape, their effective population size will unlikely be below the ∼500 where we see single population drift losses). Then, when in a meta-population landscape some populations are lost, drift will increase but most importantly the whole system will tend towards a future lower genetic diversity equilibrium. Population genetics tells us that for populations to reach equilibrium about 4Ne generations. Assuming different starting *N*_0_ values and using the information of time interval, we can then compute

**Figure S 24:**
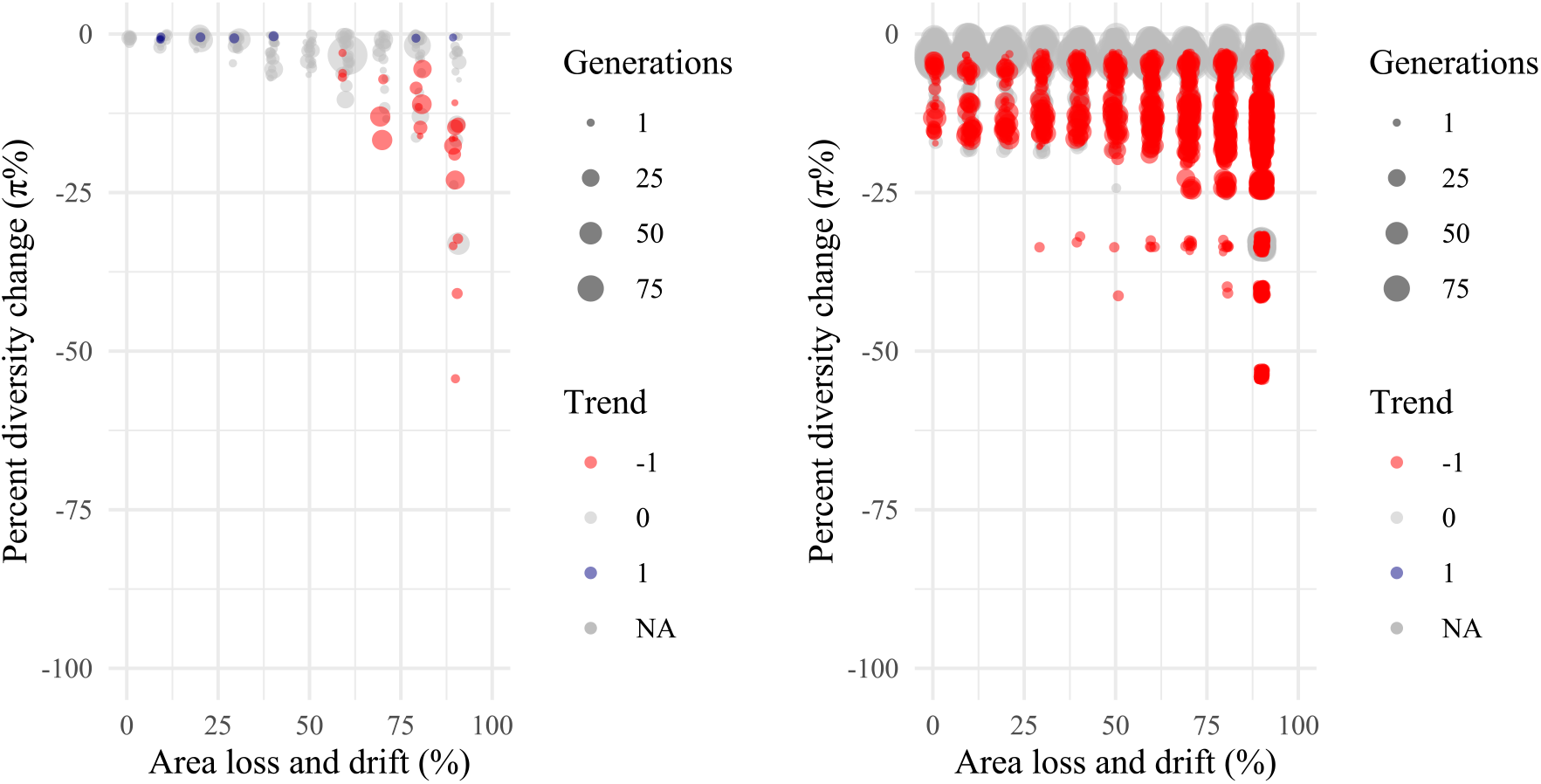
Expected immediate geographic area loss given observed π%.

#### *Average genetic diversity* π Summary

*No noise*

Median = NA% of original range lost.

Mean = NaN% of original range lost.

IQR = NA - NA% of original range lost.

Ranges = Inf - −Inf% of original range lost.

*Noise*

Median = 90% of original range lost.

Mean = 73% of original range lost.

IQR = 60 - 90% of original range lost.

Ranges = 0 - 90% of original range lost.

***Allelic richness*** *M* We have no model for this.

### Text S5: Interpret increase in genetic diversity

There are three options to interpret population genetic diversity: (1) that genetic diversity is indeed increasing species-wide, (2) genetic diversity is inflated due to changes in meta-population dynamics, (3) measurement error.

#### SINGLE POPULATION GROWTH OVER TIME – *gain scenario 1b*

**Figure S 25:**
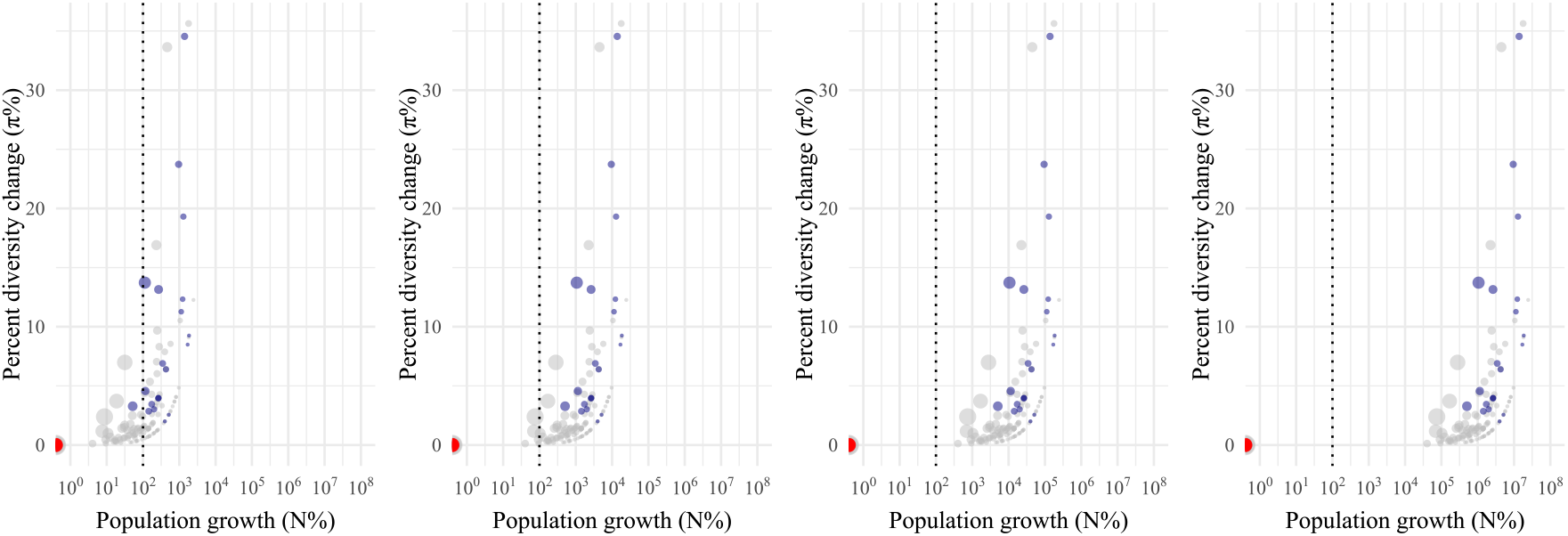
Expected population growth given increases in π%.

Fig. 27 shows plots assuming final population sizes: *N* 1 = 100, 1000, 10000, 1000000. All data points of decline area assumed to be 0%. Dotted line marks the 100% increase, or doubling of population.

Fig. 26 explains why growth of genetic diversity is even less expected than declines. It assumes a population started at 10,000 individuals, and went down to 1,000 or increased to 15,000.

**Figure S 26:**
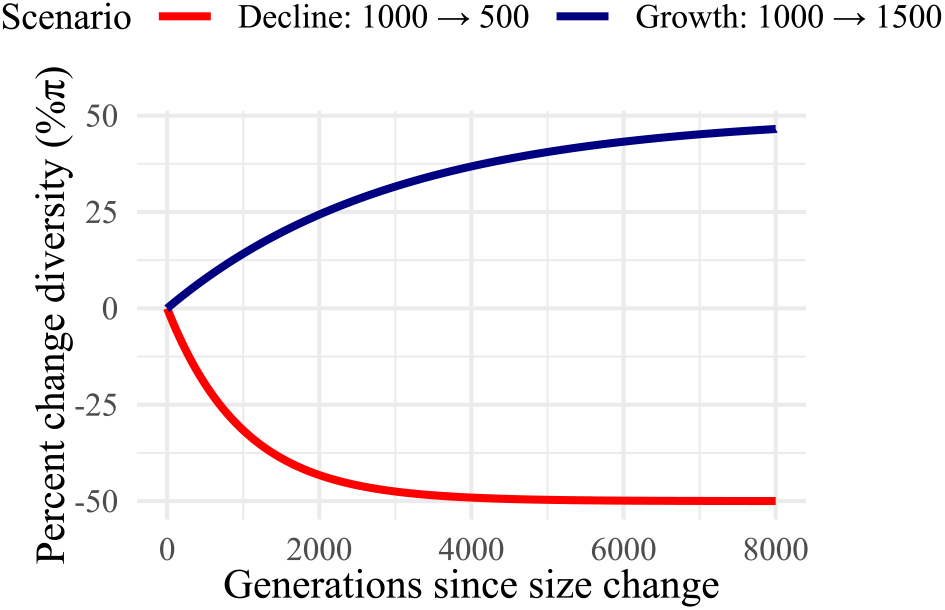
The shape of genetic diversity increases is slower than decreases.

#### SPATIAL METAPOPULATIONS MIXING – *gain scenario 2a*

The median *F*_*ST*_ required for a mixing of two popualtions to cause the percentage genetic change increase observed in positive cases will be: NA (27).

**Figure S 27:**
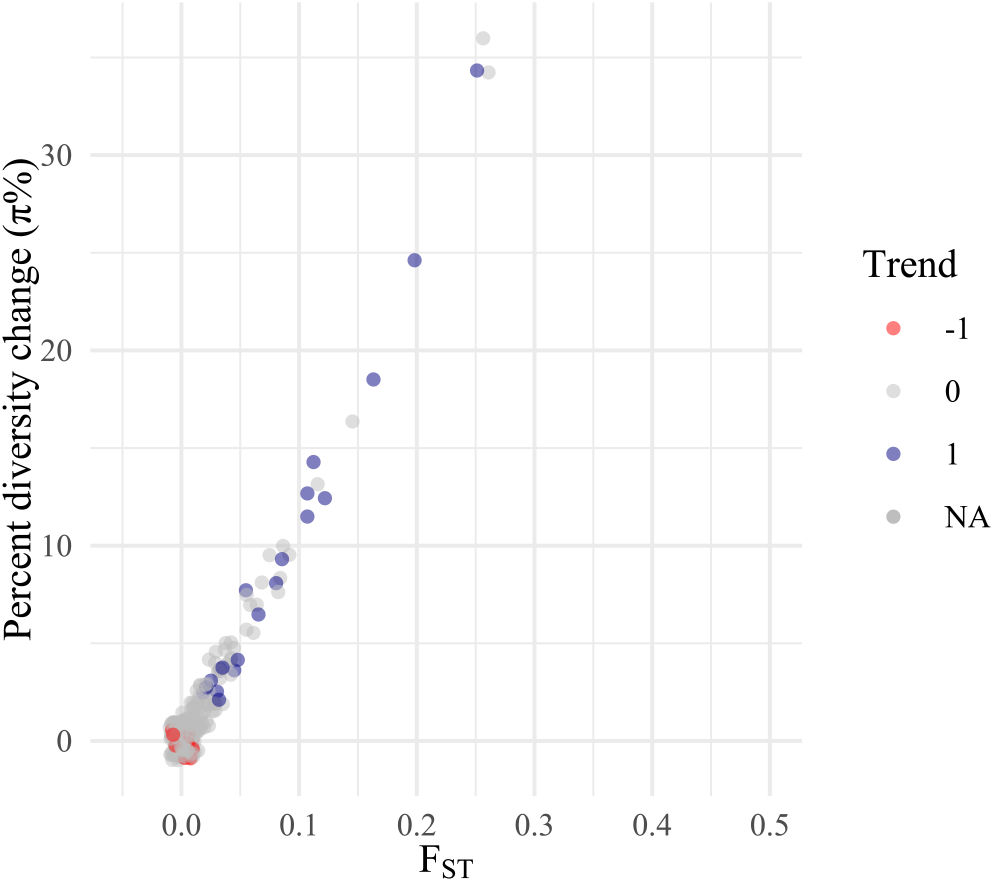
Expected *F*_*ST*_ to cause a inflation of π%.

NB 28. Plot truncated to up to +100% in genetic diversity. Negative diversity trends truncated.

This also shows that it could be that even if genetic diversity increases, area loss and fragmentation could have happened.

Expected instantaneous area contraction percentage for studies with significan decline of π :

Mean = 43% of original range.

Median = 40% of original range.

IQR = 28 - 70% of original range.

Range = 0 - 90% of original range.

### Text S6: Simple population contraction averages from LPI

As a though experiment – and to avoid controversial discussion on how or whether to measure abundance changes from LPI data – I simply take the earliest and latest census value for each LPI tracked population and compute a arithmetic average if a species has multiple observations. This yields −54% [IQR = −27 - 81%]. The median number of populations is: 2 populations [IQR = 1 - 5]. The median number of years in between earliest and latest observation is: 9.8 years [IQR = 4.4 - 17 years].

A thought experiment back-calculation of genetic diversity loss can be created using, for instance, scenario 2a, *X*_π_ = 16% [IQR 1.6 - 8.1%].

**Figure S 28:**
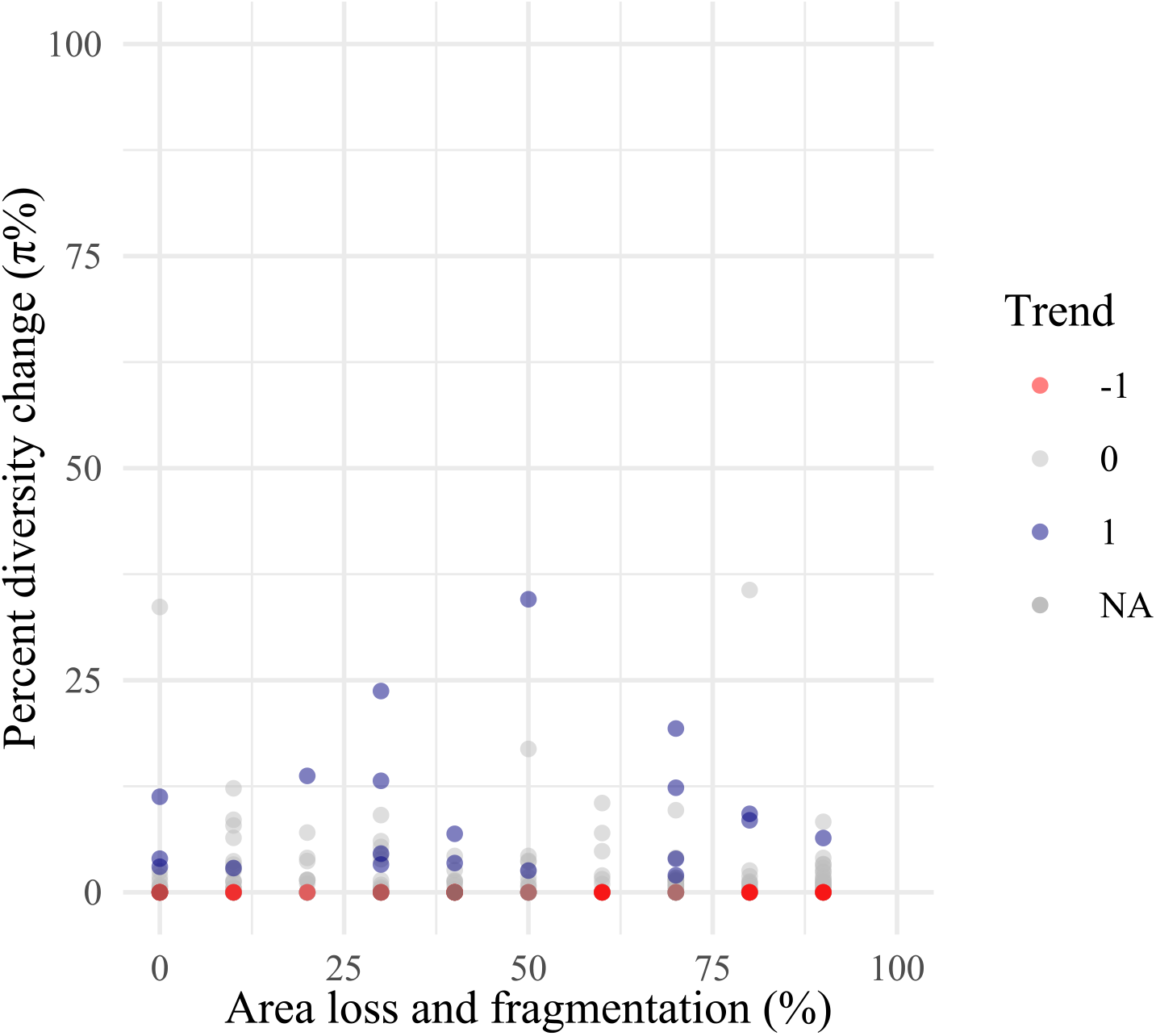
Expected noisy inflation of π% from area loss and fragmentation.

### Text S7: Limitations, caveats, and thoughts forward

#### Thoughts on data limitations

- Published studies may have too short timelines to detect signals. About 10 years or 2 generations on average are in evolutionary trends effectively instantaneous and even if populations are undergoing bottlenecks and drift changes are expected to be small. Possibly applying historical/ancient DNA technologies would enable the temporal breadth backwards (see also next point). This will help avoiding the “baseline shifting problem” in biodiversity, where our reference point of what “original diversity” was shifts constantly leading to having a recent past (already eroded nature) as our pristine nature reference.
- A problem in genomic monitoring that we are facing now and will continue happening in the future is that genetic technologies are moving fast. Therefore, comparing observations published in the literature 10 years ago is today considered obsolete genetic technologies, so meta-analysis will not make use of the best current data. This is happening in the Shaw dataset, a typical molecular ecology study measuring genetic composition of a population may include 10 microsatellites for genotyping. Nowadays, even moderate budget projects could possibly use genomic technologies to generate much more accurate values of genetic diversity (NB the measurement error rate scales 1/number of bp sequenced). This can be done even using pool sequencing (i.e. sequencing multiple individuals’ DNA in one tube) since one aims to get a population-level diversity estimate. In addition, while many genomic approaches need of assembling a genome blueprint for a species, Kmer-based approaches (digesting sequencing reads directly in DNA sequence tables) has proven to be efficient at characterizing genetic diversity (Roberts & Josephs et al. 2025, Evolution Letters).
- A problem in genetic data often ignored or not reported is the scale of measurement of genetic diversity. While population genetics classically describes the individual, the population, and the species-which has been adopted to essential biodiversity variables (Hoban et al. 2022 Biological Reviews)-these three levels are not well captured in sampling protocols nor in policy texts (CBD). The difficulty is that genetic diversity within a population may change while it not changing at the species-level scale. Likewise genetic diversity within an individual (heterozygosity or inbreeding coefficient) may change but diversity at the population is maintained. Different demographic processes, evolutionary processes, may affect these without them being problematic, in fact they may go in different directions. It seems all levels are relevant (intuitively perhaps within-species may be the most relevant) and analyses should be conducted at all such three scales.

#### Thoughts on inverse model limitations

Apart from data limitations, the inverse modeling proposed in the text also has limitations and possible improvements. No model is going to be perfect and one needs to balance usability (e.g. back of the envelope predictions vs complex process-based simulations). Here I mostly opted for back of the envelope mathematical predictions such as single population dynamics or genetic diversity-area relationships as these are very powerful to scale to many species (scenario 1a,1b,2a). For more accurate predictions and dynamics I also included some simulation/numeric based approaches (scenario 2b, Mualim et al 2024 bioRxiv) which can be extended to any population geographic conformation wanted, starting population size, migration rates, but these require further information per species.

- Shifting baseline problem underestimates impacts: Many species have already decreased in population sizes and thus contemporary measurements used in modeling (bottlenecks - to - genetics) or inverse modeling (genetic - to - bottlenecks) equations will underestimate past declines of genetic diversity before either genetic monitoring or popualtion monitoring were established.
- Survival bias underestimates impacts: by definition genetic monitoring is only going to capture genetic diversity within a population, but cannot quantify genetic diversity loss of unsampled or extinct populations. It is likely that many populations within species are being altered with habitat losses (Exposito-Alonso 2022 Science) and thus genetic diversity loss should be larger than, e.g., what is inferred in Fig. 2.
- Wild populations often do not follow Wright-Fisher evolutionary assumptions, modeling may under- or over-estimate impacts. The inferences based on evolutionary theory equations, equilibrium, sampling, necessarily use assumptions. Some assumptions involve no natural selection, random outcrossing of individuals (scenario 1a/1b), equal reproduction, etc. Underestimation may occur if, for instance, bottlenecks are impacting populations’ Ne to a larger degree due to selfing or other. Models may overestimate impacts of bottlenecks, for instance, when in nature populations have overlapping generations that maintain genetic diversity.
- Unknown refugia/reservoirs of populations containing much of a species genetic diversity will protect it to impacts elewhere thus models may over-estimate genetic diversity loss.
- Historical trends have legacy trajectories in genetic diversity. Species are not static, and are often beem changing in population size for long periods of time. Whether these are upward or downward, will have effects in genetic diversity trajectories in the future (even with contemporary rapid changes in population).

## Mathematical appendix | Response to Shaw et al. (2025)

Moi Exposito-Alonso

Department of Integrative Biology, University of California Berkeley, California, USA Howard Hughes Medical Institute, University of California Berkeley, California, USA

## SETUP

### Overview of evolutionary population genetics parameters

#### Diversity parameters

- π = nucleotide diversity or expected heterozygosity (identical under biallelic loci)
- *S* = allelic richness, segregating sites, number of variable positions, or number of mutations in a DNA strand. **Note**: We use *M* instead of *S* to avoid confusion with species richness from ecology, and avoid *A* (alleles) to prevent confusion with area of habitat
- θ = diversity parameter of a population; under equilibrium θ = 4*N*_*e*_*μ*

#### Population parameters

- *N*_0_ = population size at some past time (long-term equilibrium size)
- *N*_1_ = population size after instantaneous reduction from human impact or contraction (smaller population, not in equilibrium)
- *N*_*t*_ = population size at time *t*
- *A*_0_ = original habitat area of a species with multiple populations
- *A*_1_ = reduced habitat area after human impact or contraction

#### Parameters of change

- *X*_π_ = fraction loss of genetic diversity π (intuitive % loss when *X*_π_ × 100)
- *X*_*M*_ = fraction loss of allelic richness *M*
- *X*_*N*_ = fraction loss of population size = 1 − (*N*_1_/*N*_0_)

#### Stable state of genetic diversity in a population

Under equilibrium conditions (where *N*_*e*_ = *N*_0_), the expected genetic diversity follows from the coalescent process. The total length of all coalescent branches in a population genealogy is *L*_total_, and the number of mutations is *M* = *μL*_total_.

The expected genealogy size under the coalescent is:

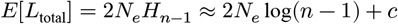

where: - 2*N*_*e*_ = average time to coalescence between any two samples - *H*_*n*−1_ = harmonic number representing progressive coalescent of all samples - This can be approximated with log(*n* − 1) for large *n*

Therefore:

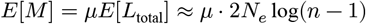

For nucleotide diversity π (equivalent to expected heterozygosity under biallelic variants), under equilibrium:

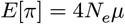

This represents the average number of genetic variants between two samples along a DNA stretch, derived from the average pairwise coalescent time:

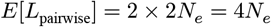

### Key insight

*M* is more sensitive to low-frequency variants (any mutation in *≥*1 individual counts), while π is more sensitive to intermediate-frequency variants. For a biallelic variant with frequency *p*, the probability that two random samples differ is 2*p*(1 − *p*), maximized at *p* = 0.5.

#### Summary of loss and gain scenarios

We examine genetic diversity changes for both π and *M* under different spatio-temporal scenarios:

**Spatial scenarios:** - **Scenario 1**: Single population declining (applicable to threatened species) - **Scenario 2**: Multiple populations with fraction of populations lost

**Temporal scenarios:** - **Scenario a**: Sudden immediate decline (direct mortality effect) - **Scenario b**: Loss due to demographic stochasticity over time (genetic drift)

**Combined scenarios**: 1a, 1b, 2a, 2b

### DIVERSITY DECLINES

#### Loss scenario 1a: Immediate population contraction (no drift effects)

When a population contracts immediately from *N*_0_ to *N*_1_ within a single generation, we can model this as statistical subsampling.

**Loss of π** For nucleotide diversity, the sampling effect creates a downward bias. Under the coalescent, when sampling *n* individuals from a finite population of size *N*, there’s a probability 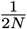 that samples coalesce to the same ancestor (yielding zero diversity). Additionally, finite sampling creates bias captured by 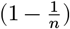.

The relationship becomes:

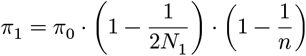

**Key result**: Loss is proportional to 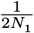, so 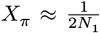.This means: - Very small genetic diversity loss unless *N*_1_ < 40 - Even at *N*_1_ = 40: *X*_π_ = 0.0125 (1.25% loss) - π is largely unaffected by bottlenecks within one generation unless near-complete population eradication occurs

**Loss of alleles M** Since *M*_0_ ∝ log(*N*_0_) under equilibrium, the fractional loss is:

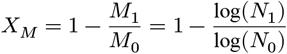

Substituting *N*_1_ = *N*_0_(1 − *X*_*N*_):

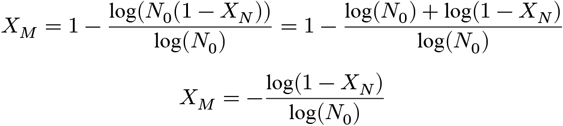

**Key insights**: - Genetic diversity loss *X*_*M*_ is log-proportional to population size loss log(1 − *X*_*N*_) - Species with larger initial populations lose smaller fractions of genetic diversity - **Example**: *N*_0_ = 10, 000, 50% reduction (*X*_*N*_ = 0.5) → *X*_*M*_ = 7.5 allelic diversity loss

#### Loss scenario 1b: Effect of drift over time

**Loss of π over time** Changes in nucleotide diversity over time follow the classic formula:

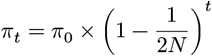

The fractional loss after *t* generations:

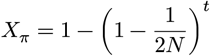

Rearranging to solve for effective population size:

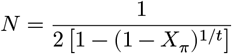

**Example**: For *N* = 100 individuals: - *X*_π_ = 0.5% per generation - ∼5% loss over 10 generations - This insight supports the conservation genetics recommendation of maintaining *N* > 500

**Loss of alleles M over time** The probability that an allele with frequency *p*_*i*_ is lost in one generation is (1 − *p*_*i*_)^2*N*^.

The allelic richness after one generation:

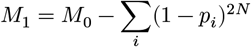

**Simplified approximation**: Discretizing frequency distribution into: - Rare alleles (<1%): high loss probability (>50%) - Uncommon alleles (1-10%): moderate loss probability

- Common alleles (>10%): very low loss probability (<0.1%) Using the standard frequency distribution *f* (*p*) ∝ θ/*p*:

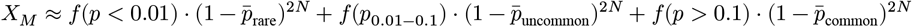

**Example**: 90% population decline (*N*_0_ = 1000 → *N*_1_ = 100): - *X*_*M*_ ≈ 15% allelic richness loss per generation Over *t* generations, survival probability for allele with frequency *p*:

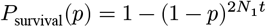

**Loss scenario 2a: Spatial geographic contraction (immediate)**

**Loss of π with population structure** We use the fixation index *F*_*ST*_ (Nei’s definition):

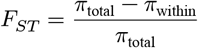

where π_total_ = π_within_ + π_between_.

For a species with *k* populations, total genetic diversity is:

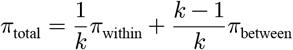

We can express components in terms of *F*_*ST*_ :

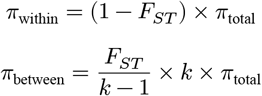

After losing *x* populations, the new total diversity:

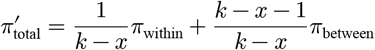

The fractional change:

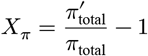

Substituting and simplifying:

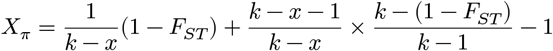

**Key insights**: - Minimum populations (*k* = 2): *X*_π_ = *F*_*ST*_ - As *k* increases, loss per population decreases -

**Example**: *F*_*ST*_ = 0.2, 50% area contraction → *X*_π_ ≈ 0.2% to 20% (depending on *k* = 2 to 100)

**Continuous space approximation**: Using the Genetic Diversity-Area Relationship (GDAR):

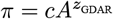

where empirically *z*_GDAR_ ≈ 0.05. This gives:

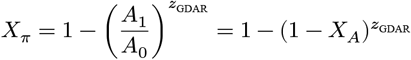

**Example**: 50% area reduction → *X*_π_ ≈ 3.4%

**Loss of alleles M with spatial contraction** Using the Mutations-Area Relationship (MAR):

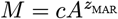

where empirically *z*_MAR_ ≈ 0.25. This gives:

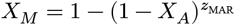

**Example**: 50% area reduction → *X*_*M*_ ≈ 19% allelic richness loss

#### Loss scenario 2b: Spatial contraction with drift over time

This scenario combines spatial extinction with ongoing demographic stochasticity. The temporal dynamics require sophisticated numerical methods (e.g., Wright-Fisher moments) and are beyond simple analytical solutions. These models are particularly relevant for:

- Non-threatened species with moderate to large geographic ranges
- Global within-species diversity loss (vs. classic conservation studies of extremely reduced populations)
- Long-term predictions of genetic diversity under habitat fragmentation

### DIVERSITY INCREASES

#### Gain scenario 1b: Population growth over time

Starting from the classic equation, rearranged for expansion. π_0_ = 4*N*_0_*μ* (diversity before expansion). π_∞_ = 4*N*_1_*μ* (new equilibrium after expansion). π_*t*_ = diversity *t* generations after expansion The trajectory follows:

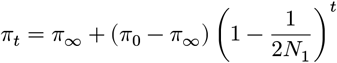

#### Derivation of fractional change

Define the population growth ratio: 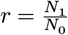

Then 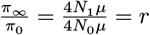

The fractional change from baseline:

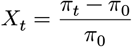

Substituting and simplifying:

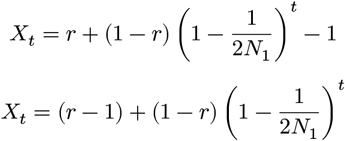

Expressing in terms of population change 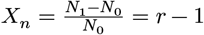:

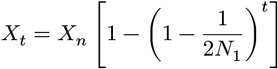

Or in terms of original population size:

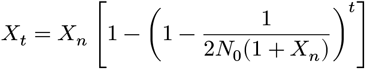

**Interpretation**: The genetic diversity approaches the new equilibrium value exponentially, with rate depending on the expanded population size *N*_1_.

#### Gain scenario 2a: Spatial expansion (*F*_*ST*_ based)

When populations expand spatially and come into contact, genetic diversity increases. The proportional increase can be derived from population differentiation, starting from: 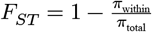.

The relationship between diversity gain and population structure can be re-arranged then as:

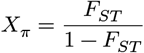

Conversely: 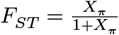.

**Interpretation**: Higher population differentiation leads to greater potential diversity gains upon population mixing or expansion.

